# Bayesian Cooperative Learning for Multimodal Integration

**DOI:** 10.1101/2025.10.23.684056

**Authors:** Saptarshi Roy, Sreya Sarkar, Erina Paul, Piyali Basak, Nengjun Yi, Himel Mallick

## Abstract

Bayesian multimodal models that integrate data from multiple sources, studies, or modalities have garnered considerable attention in recent years. However, existing methods either rely on computationally expensive Markov chain Monte Carlo (MCMC) schemes or adopt variational approaches that forgo principled uncertainty quantification. To address this limitation and cater to practical needs, we abandon the MCMC framework and turn to resampling-based posterior inference for multimodal integration. Our method, Bayesian Cooperative Learning (BayesCOOP), embeds fast maximum a posteriori (MAP) estimation within a Bayesian bootstrap, combining a novel jittered group spike-and-slab prior with an efficient expectation–maximization (E M) coordinate descent algorithm under randomly weighted data perturbations. Averaging posterior summaries (MAP estimates) across bootstrap replicates yields approximate posterior samples that retain Bayesian interpretability while avoiding the computational burden of traditional sampling-based inference. We establish theoretical connections between BayesCOOP’s pseudo-posterior averaging and posterior contraction principles, demonstrating near-optimal posterior consistency under sparsity. Extensive simulation studies and analyses of pregnancy multi-omics datasets demonstrate that BayesCOOP substantially outperforms state-of-the-art early, intermediate, and late fusion approaches in estimation, prediction, and uncertainty assessment. The open-source implementation of BayesCOOP is available at https://github.com/himelmallick/BayesCOOP.

## 1. Introduction

Modern biomedical research increasingly relies on integrating heterogeneous data measured across multiple molecular, imaging, and clinical modalities to understand complex biological systems and improve outcome prediction. These multimodal datasets, which often capture complementary biological processes, hold great promise for uncovering latent mechanisms of health and disease. Yet, integrating them poses formidable statistical and computational challenges due to their high dimensionality, varying measurement scales, and missingness patterns (Xu and McCord, 2022). Bayesian data integration provides a principled framework for tackling such challenges by coherently combining prior biological knowledge with data-driven learning while enabling probabilistic uncertainty quantification (Mallick et al., 2024; Anceschi et al., 2024). However, despite its theoretical appeal, fully Bayesian inference via Markov chain Monte Carlo (MCMC) can be computationally prohibitive and unstable in large-scale multimodal settings. Moreover, MCMC-based posteriors can exhibit high Monte Carlo variability, leading to results that are difficult to reproduce across runs or tuning configurations (Huggins and Miller, 2024). Variational Bayesian methods offer a computationally efficient alternative (Liu and Zhong, 2024, 2025) but often underestimate posterior uncertainty due to restrictive factorization assumptions and can be sensitive to model misspecification (Srivastava et al., 2018; Mauri and Dunson, 2025). These limitations have spurred growing interest in scalable Bayesian approaches that retain interpretability and improve both efficiency and reproducibility (Huggins and Miller, 2024; Nie and Ročková, 2023).

One area where these challenges are particularly evident is maternal–fetal health research, which increasingly leverages multimodal biological measurements to understand the determinants of pregnancy outcomes (Ghaemi et al., 2019; Stelzer et al., 2021). For instance, recent studies have collected multi-omics and immunologic data–such as cell-free transcriptomics, cytokine panels, microbiome profiles from multiple body sites, and plasma metabolomics and proteomics—from the same participants to characterize the interplay between immune, metabolic, and microbial processes during pregnancy (Ghaemi et al., 2019; Stelzer et al., 2021). Each modality offers unique prognostic information, yet they are biologically interconnected. For example, microbial composition can influence immune and metabolic pathways, which in turn affect cytokine levels and gene expression relevant to maternal adaptation. These complex dependencies raise several modeling questions: Should all data layers be integrated for prediction or only a subset? How can shared biological signals be amplified while suppressing modality-specific noise? And can scalable bi-level feature selection identify both key modalities (group-level sparsity) and the most predictive features within them (within-group sparsity)?

To tackle these challenges, several methods have been proposed to integrate multiple data layers for improved prediction and interpretation, encompassing both Bayesian and frequentist paradigms. At one end of the spectrum, Bayesian factor models learn shared and modality-specific latent factors that explain cross-modality dependence; these models are interpretable but rely on strong distributional assumptions and computationally intensive inference (De Vito et al., 2021; Grabski et al., 2023; Mauri and Dunson, 2025; Anceschi et al., 2024; Samorodnitsky et al., 2024). Variational adaptations of these models have achieved notable computational gains (Hansen et al., 2022, 2025; Liu and Zhong, 2024, 2025), yet such approximations tend to underestimate posterior uncertainty and exhibit sensitivity to model misspecification (Srivastava et al., 2018; Mauri and Dunson, 2025). At the other end, non-factor-analytic approaches either (i) use late fusion schemes that aggregate modality-specific predictors at the outcome level and therefore ignore cross-modality dependence (Stelzer et al., 2021; Ghaemi et al., 2019; Mallick et al., 2024), or (ii) adopt dependence-aware frameworks that model cross-view dependency without introducing latent factors, typically within a frequentist paradigm (Ding et al., 2022). While such frequentist approaches are computationally efficient and enable multiview feature selection (the simultaneous identification of informative modalities and features within them), they lack a coherent Bayesian mechanism for posterior inference or uncertainty assessment. As a result, existing approaches either offer comprehensive uncertainty quantification at prohibitive computational cost or enable scalable multiview feature selection with little guidance on uncertainty, leaving a critical gap in achieving scalable and simultaneous uncertainty quantification and feature selection for multimodal integration.

To address the practical need and methodological void, we develop Bayesian Cooperative Learning (BayesCOOP), a scalable Bayesian framework for supervised multimodal integration that embeds fast maximum a posteriori (MAP) estimation within a Bayesian bootstrap (Rubin, 1981). Unlike existing Bayesian multimodal methods, BayesCOOP rethinks uncertainty quantification through a resampling-based Bayesian perspective, in which repeated deterministic optimization under randomly weighted data realizations yields reproducible approximations to posterior variability. By replacing stochastic sampling and restrictive variational assumptions with deterministic optimization within a Bayesian bootstrap, BayesCOOP achieves stable, interpretable, and computationally efficient inference in high dimensions. Further, BayesCOOP employs a novel *jittered group spike-and-slab double exponential prior* to achieve bi-level sparsity, while averaging posterior summaries across bootstrap replicates to form a pseudo-posterior distribution. Through numerical experiments, we demonstrate that BayesCOOP significantly surpasses existing Bayesian and frequentist methods in terms of both estimation and prediction.

The subsequent sections of the paper are structured as follows. In Section 2, we describe the BayesCOOP method and establish theoretical guarantees of the proposed methodology. To assess the performance of BayesCOOP, simulation studies are conducted in Section 3, and real data analyses are presented in Section 4. In Section 5, we briefly discuss potential avenues for further research in this field. Technical proofs and additional materials are provided in the **Supplementary Materials**.

## 2. Bayesian Cooperative Learning for Multimodal Integration

In this section, we first introduce the notational framework for supervised multiview learning (Section 2.1). Following prior work (Ding et al., 2022), we formulate the multiview problem as an intermediate fusion framework that encourages agreement among predictions across views while retaining view-specific effects through an agreement parameter. This formulation allows us to devise a suitable data augmentation strategy that reformulates the multiview likelihood into a more tractable augmented likelihood. In Section 2.2, we present the Bayesian bootstrap methodology incorporating a novel jittered group spike-and-slab Laplace prior, within which we embed a computationally efficient expectation–maximization (EM)-based coordinate descent algorithm for MAP estimation. In Section 2.3, we outline the theoretical underpinnings of our proposed framework and specify the conditions under which posterior consistency of the pseudo-posterior is achieved. Figure 1 summarizes our end-to-end workflow and implementation, available as an open-source R package at https://github.com/himelmallick/BayesCOOP.

**Figure 1.**
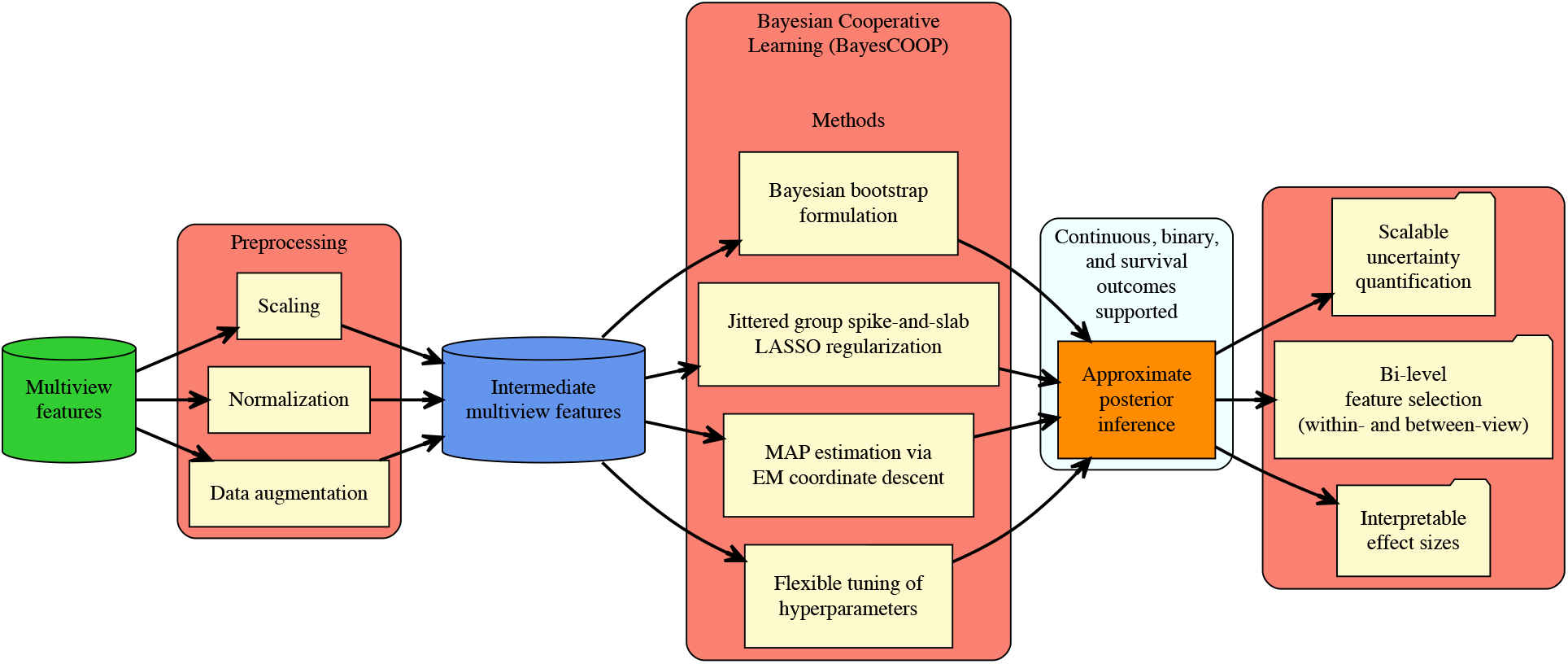
Bayesian Cooperative Learning for Multimodal Integration. Starting with quality-controlled multiview data (normalized and scaled), BayesCOOP performs a data augmentation step to transform the data into an intermediate, dependence-aware representation, corresponding to the intermediate fusion strategy described in **Section 2.** The modeling framework comprises (1) a Bayesian bootstrap formulation, (2) a *jittered* group spike-and-slab double exponential prior, and (3) a computationally efficient EM coordinate descent algorithm for MAP estimation. In addition to scalable bi-level feature selection that identifies multiview features with interpretable effect sizes for downstream analysis, BayesCOOP provides scalable uncertainty quantification through approximate posterior sampling. The implementation is modular, allowing multiple customization options for practical applications.

### 2.1 Prior Elicitation

To achieve the generality of our approach, we first set up the model for multiview learning. Consider a multiview dataset with *n* data points and *d* data views. Let ***X***^(1)^, ***X***^(2)^, …, ***X***^(*d*)^ be the feature matrices corresponding to *d* views, where 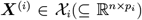, and let ***y*** = (*y*_1_, …, *y*_*n*_)^⊤^ ∈ 𝒴 denote the response vector. We seek to learn a map *g* : 𝒳 → 𝒴, where 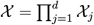, that combines information from multiple views to improve prediction accuracy of ***y***. Within a linear regression framework, this reduces to estimating the parameter vector 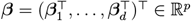, where 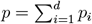, under the model

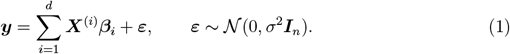

In most practical applications, each view is high-dimensional or ultra-high-dimensional, where the number of variables *p*_*i*_ far exceeds the sample size *n*. Specifically, we consider settings where 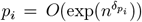 for some positive constants *δ*_*pi*_. We assume an inherent grouping structure among predictors, as is typical in multiview datasets, and that the view-specific regression coefficients are doubly sparse meaning that only some of the groups are active in driving the outcome, and for each active group, only a small subset of variables significantly impacts the outcome. Recognizing that efficient bi-level variable selection is essential for accurate estimation and interpretability in complex, high-dimensional multiview settings, we introduce a novel multiview shrinkage prior, the group spike-and-slab Laplace prior, as delineated in the sections that follow.

Before introducing the multiview shrinkage prior, we first define the *regularized multiview likelihood function*, which introduces an agreement parameter that penalizes discrepancies between predicted responses across pairs of views (Ding et al., 2022):

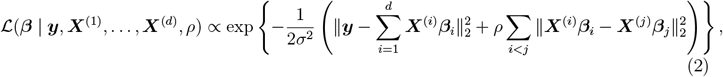

where *ρ >* 0 controls the degree of information sharing across views. This formulation simplifies to early fusion and late fusion when *ρ* = 0 and *ρ* = 1, respectively. This construction penalizes large deviations between fitted responses from different modalities, thus encouraging cross-view coherence while preserving view-specific effects.

A key computational insight is that the cross-view agreement term in (2) admits a *data augmentation representation*. Specifically, the agreement term can be expressed as additional pseudo-observations appended to the data, yielding an augmented response vector.

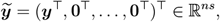

With 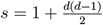, and an augmented design matrix 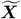 of the form

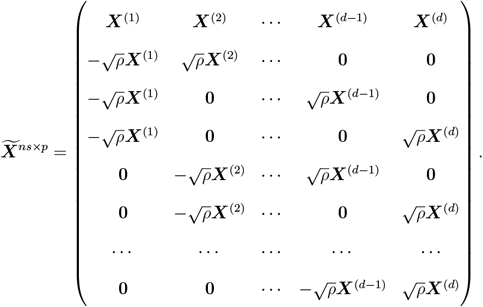

Under this construction, the likelihood in (2) can be equivalently rewritten as a standard Gaussian likelihood on the augmented data:

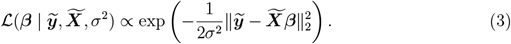

Let the coefficients be clustered into *d* informative groups, *C*_1_, *C*_2_, …, *C*_*d*_. We consider a hierarchical group spike-and-slab double exponential prior as follows:

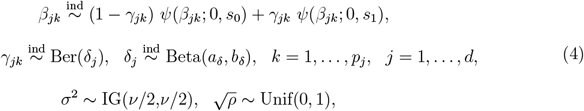

where *ψ*(*·*; *a, b*) denotes the Laplace density with location *a* and scale *b*. The shrinkage level is controlled via *s*_1_ ≫ *s*_0_ *>* 0, so that when *γ*_*jk*_ = 1, the (*j, k*)-th coefficient has a diffuse prior, and otherwise a sharply concentrated prior near zero. The group-level inclusion probability *δ*_*j*_ enables adaptive shrinkage across views, facilitating structured multiview regularization.

A straightforward Gibbs sampler can be implemented by expressing the double exponential prior as a scale mixture of Gaussians with an exponential mixing distribution (Ročková and George, 2018; George and McCulloch, 1993). This reparameterization yields conditional conjugacy under the multiview group spike-and-slab specification.

### 2.2 Pseudo-Posterior Sampling with Embedded MAP Estimation

Although such a full MCMC approach offers rich uncertainty quantification, it becomes computationally expensive in high-dimensional settings, especially as the parameter space scales with the number of views. To circumvent these challenges while preserving the benefits of the Bayesian inferential framework, we deploy the jittered Bayesian bootstrap strategy (Nie and Ročková, 2023) for efficient approximate posterior sampling of the ultra-high-dimensional regression coefficient ***β***. This method bypasses MCMC sampling by (1) independently optimizing randomly perturbed, reweighted likelihood functions and (2) randomly recentering the prior mean of the individual coordinates of ***β***. This dual-perturbation approach is advantageous in many ways. First, the approximate posterior sampling scheme is founded upon MAP estimation of ***β*** based on a reweighted likelihood, which enjoys the computational scalability of optimization over Gibbs sampling from a full conditional distribution. Second, with suitably chosen distributions of the perturbing weights, the resultant pseudo-posterior is theoretically guaranteed to concentrate around the true posterior despite avoiding actual MCMC sampling (Section 2.3). Third, recentering the prior around randomly jittered locations prevents excessive posterior concentration at zero for weak or inactive signals, addressing a key limitation of naive MAP-based approaches that often underestimate uncertainty. We thus formalize the hierarchical model with a jittered group spike-and-slab prior for our method as follows.

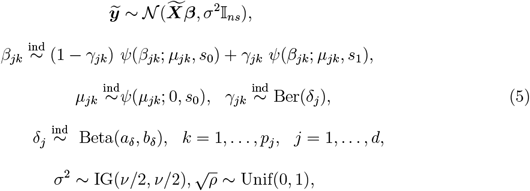

where we set *a*_*δ*_ = *b*_*δ*_ = 1 as default choices. The parameter *ν* of the inverse gamma distribution is set to 1 to make it relatively non-influential.

To conduct approximate posterior inference for BayesCOOP via embedded MAP estimation, we develop an efficient algorithm that integrates an Expectation–Maximization (EM) procedure based on a reweighted likelihood with an extremely fast cyclic coordinate descent algorithm. Although similar optimization-based approaches have appeared in single-view settings (Ročková and George, 2018; Tang et al., 2018), none have been formulated for multiview models within the Bayesian bootstrap framework.

In particular, at any *t*-th iteration, we first sample the agreement parameter 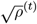 from its full conditional distribution. Notably, this represents a key advance over the frequentist Cooperative Learning framework (Ding et al., 2022), which requires computationally intensive cross-validation to estimate the corresponding agreement parameter. Subsequently, given *ρ*^(*t*)^, we sample the weight 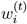 corresponding to the *i*-th observation in the augmented setup (*i* = 1, …, *ns*). Specifically, as recommended by Nie and Ročková (2023) for parametric models, we draw 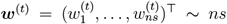 Dirichlet(1, …, 1), and obtain the MAP estimator 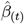 by maximizing the jittered pseudo-posterior as below

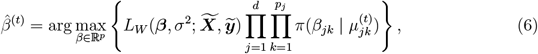

where 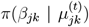 denotes the jittered group spike-and-slab Laplace prior as described in (5), and the reweighted likelihood is

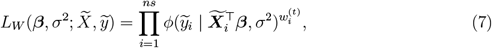

with *ϕ*(*·*; *µ, σ*^2^) representing the Gaussian density with parameters *µ* and *σ*^2^. Finally, we complete the iteration by sampling *σ*^2(*t*)^ from its full conditional distribution.

The full procedure, which embeds MAP estimation within the Bayesian bootstrap framework, is outlined in **Algorithm 1**. For brevity, the mathematical details and corresponding pseudocode illustrating the extension of BayesCOOP to generalized outcomes are deferred to the **Supplementary Materials**. Having outlined the algorithmic and modeling extensions, we next turn to the theoretical properties of BayesCOOP.

### 2.3 Theoretical Guarantees

Recent advances in scalable Bayes have highlighted the promise of resampling-based posterior approximation methods (Huggins and Miller, 2024; Nie and Ročková, 2023). For instance, Nie and Ročková (2023) introduced a compelling paradigm in which fast MAP optimization is performed on jittered priors and randomly weighted datasets to approximate the full posterior distribution. Complementarily, Huggins and Miller (2024) proposed the bagged posterior framework, which averages posterior distributions over bootstrapped datasets to enhance reproducibility and uncertainty calibration under model misspecification. Collectively, these works demonstrate that such pseudo-posteriors can retain the desirable contraction properties of the exact Bayesian posterior in high-dimensional regimes, achieving near-minimax rates for both sparse mean estimation and regression problems.

Building upon these insights, we investigate analogous theoretical properties for our proposed BayesCOOP framework. In the sequel, we present two key results: (i) a recovery guarantee for the support size of the posterior mode under our hierarchical spike-and-slab prior with group-specific shrinkage and (ii) a rate-optimal *ℓ*_2_-contraction result for the posterior pseudo-distribution around the true sparse regression vector. These results establish that, under appropriate design conditions and prior scaling, BayesCOOP inherits the same asymptotic efficiency guarantees as Nie and Ročková (2023) while accommodating more intricate structural constraints on ***β***_0_. Next, we describe the key assumptions that provide the technical foundation for our theoretical results.

#### Algorithm 1

Pseudo-Posterior Sampling with Embedded MAP Estimation

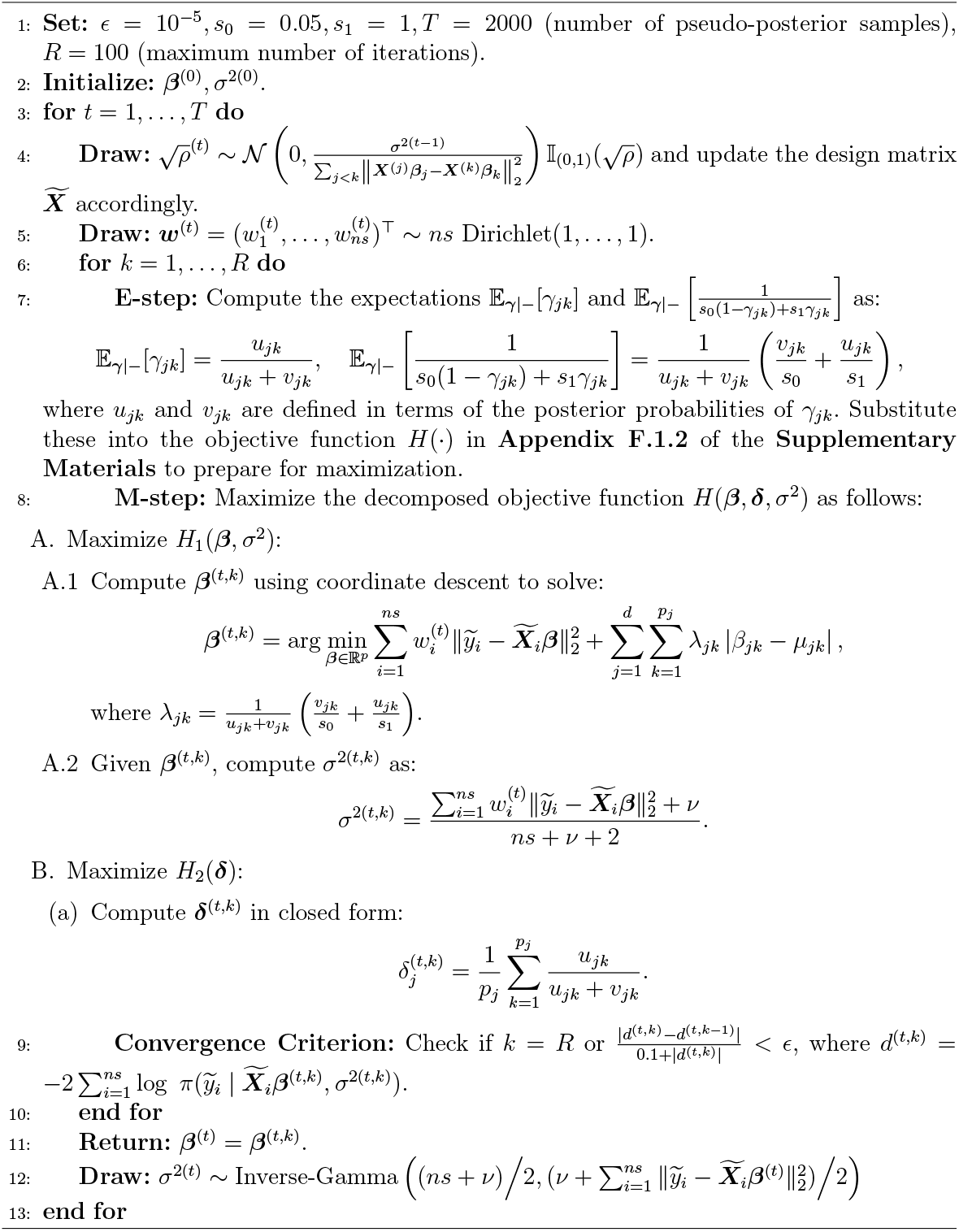

Assumption 1: Suppose the nonnegative random vector **w** = (*w*_1_, …, *w*_*ns*_)^⊤^ is drawn from a distribution *π*(**w**) satisfying the following regularity conditions:

1. Each component has unit mean, i.e., 𝔼[*w*_*i*_] = 1 for all 1 ⩽ *i* ⩽ *ns*.
2. There exists a constant *m* ∈ (0, 1) such that

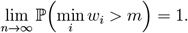
3. There exists a constant *M >* 1 such that

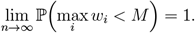
4. The marginal variances and pairwise covariances are uniformly small, satisfying

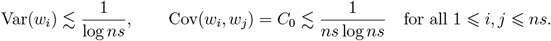

Assumption 2 (Eigenvalue condition on the design matrix): The design matrix 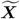 satisfies the sparse eigenvalue condition 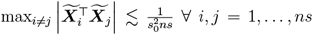, and *ξ*_0_ *>* 0 satisfies max 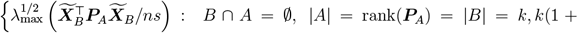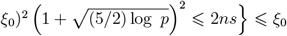. Condition 6 in Nie and Ročková (2023), pertaining to total sparsity *q* and tolerance *D <* 1, remains unchanged in our setting.

Assumption 3 (Group prior scaling): The group-specific inclusion parameters satisfy 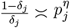, for all *j* = 1, …, *p*, with *η* ⩾ 1.

Assumption 4 (Global spike shrinkage): The global spike scale satisfies *s*_0_ ≍ *p*^−*γ*^, with *γ* ⩾ 1, and *η* + *γ >* 1.

Assumption 1 constrains the random weights to remain bounded and weakly dependent, preserving the effective sample size and preventing distortions in the weighted likelihood. Assumption 2 imposes a sparse eigenvalue condition on the design matrix, maintaining identifiability and well-conditioned geometry even in high-dimensional regimes. Assumptions 3 and 4 regulate the group-specific and global shrinkage parameters, ensuring a balance between sparsity and signal retention. Together, these conditions align the stochastic weighting, design structure, and prior scaling to guarantee consistent, well-behaved, and interpretable inference.

Theorem 1 (BayesCOOP Model Size Recovery): *Consider the linear model* 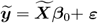 *with* ***ε*** ~ 𝒩 (0, *σ*^2^***I***_*ns*_), *where* 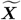 *is an ns × p design matrix with p > n, and* ***β***_0_ ∈ ℝ^*p*^ *is sparse and partitioned into d groups* 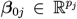 *with* 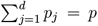 *With group inclusion probabilities δ*_*j*_ ∈ (0, 1) *and fixed variance σ*^2^, *the total sparsity is* 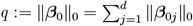 *Let* 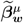 *denote the posterior mode under BayesCOOP with a jittered prior on* ***β*** *centered at* ***µ*** *and bootstrap weights* ***w***. *Assuming 1–4 hold and setting* 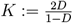, *we obtain:*

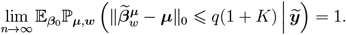

Theorem 2 (Minimax rate): *Under the same conditions as in Theorem 1, the pseudo-posterior with embedded MAP estimation concentrates at the near-minimax rate, that is*,

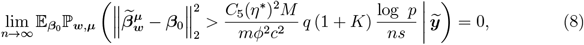

*where* 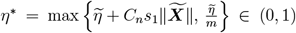,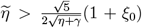 *with C*_*n*_ → ∞, *c* = *c*(*η*^∗^; ***β***), *ϕ* = *ϕ*(*C*(*η*^∗^; ***β***_0_)), *whose definitions are provided in Nie and Ročková (2023)*.

While Theorem 1 establishes that the BayesCOOP estimator consistently recovers the true model size asymptotically—selecting at most a constant multiple of the true number of active coefficients—Theorem 2 further shows that the estimator attains near-minimax convergence in *ℓ*_2_ error, ensuring optimal contraction of the pseudo-posterior around the true parameter at the rate *q*(log *p*)*/*(*ns*). The outlines of the proofs for both theorems are provided in the **Supplementary Materials**.

## 3. Simulation Studies

The increasing prevalence of multimodal studies is outpacing our ability to develop integrative methods for high-dimensional multimodal data analysis. Despite the development of numerous multimodal fusion techniques in recent years, evaluating the comparative performance of multimodal inference on realistically complex biological data remains a fundamental challenge, as neither a gold standard nor a realistic synthetic data generator exists (Mallick et al., 2024; Anceschi et al., 2024). To date, only three multiview simulators have been developed—InterSIM (Chalise et al., 2016), MOSim (Monzó et al., 2018), and OmicsSIMLA (Chung and Kang, 2019)–none of which (1) allow multivariable spike-ins necessary for evaluating multiview feature selection, (2) induce realistic feature–metadata and feature–feature correlations within and across data layers, or (3) provide flexibility to accept arbitrary user-defined template data as input to their synthetic data generation pipelines.

In the absence of such ‘gold standards’, we adopted a simulation scheme from a recent publication that integrated the InterSIM framework with arbitrary multivariable spike-ins (Mallick et al., 2024). This approach provides one of the first frameworks to characterize the properties that make true associations difficult to recover from multimodal data—or, conversely, easy to miss as false negatives. Using this method, we created deliberately challenging test cases for BayesCOOP and other published methods, offering a contrast to previous studies that relied primarily on simpler synthetic data (Ding et al., 2022). While prediction accuracy is routinely evaluated in many studies, multiview feature selection has rarely been emphasized—despite being a critical consideration for both current and future multimodal methods.

To this end, we generated multiple interrelated data types with realistic intra- and inter-relationships, based on DNA methylation, mRNA expression, and protein expression data from an ovarian cancer cohort in The Cancer Genome Atlas, using the R package *InterSIM* (Chalise et al., 2016). To ensure realism, we used the simulated multi-omics features (i.e., the **X**’s) from InterSIM to generate a continuous outcome, and conducted a series of simulation studies by varying the data-generating mechanisms, incorporating different linear and non-linear effects, and exploring a range of sparsity levels.

For each of these settings, we simulated 25 replicates of training and test datasets (*n* = 500), each with 658 features across three layers (131 gene expression, 367 methylation, and 160 protein features) and varied the sparsity level: sparse (10% non-zero), moderately sparse (30% non-zero) and dense (50% non-zero). Additional results for a smaller sample size (*n* = 200) are included in the **Supplementary Materials** (**Figs. S1-S2**). The training and test datasets are split with a fixed ratio of 70% to 30% across all replicates. The signal-to-noise ratio (SNR) is varied between low (SNR = 5) and high (SNR = 10) to regulate the level of noise in the data, where SNR is defined as 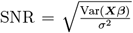. We simulate data from the true model ***y*** = *f*_0_(**X**) + ***ε, ε*** ~ N(**0**, *σ*^2^*I*), where *σ*^2^ is chosen such that the desired level of SNR is achieved. We consider the following data generation mechanisms:

**(1) Simulation 1 (“Fully linear”):** In this example, *f*_0_(**X**) = **X*β***, where only 10% of the linear coefficients are non-zero, drawn randomly from a log-normal distribution such that it leads to realistically varying log_2_-fold changes between (−3, 3) representing both modest (e.g., *<* 2-fold differences) and strong (e.g., 8-fold) effect sizes using the Splatter simulation framework (Zappia et al., 2017).

**(2) Simulation 2 (“Fully non-linear”):** In this example, *f*_0_(**X**) = 10 sin(*πX*_1_*X*_2_) + 20(*X*_3_ − 0.5)^2^ + 10*X*_4_ + 5*X*_5_, which includes non-linearity and interaction effects among five randomly chosen multi-omics features (*X*_1_, …, *X*_5_). The function *f*_0_, proposed initially by Friedman (1991), has non-linear dependence on the first three variables *X*_1_, *X*_2_ and *X*_3_, linear dependence on *X*_4_ and *X*_5_, and incorporates a non-linear interaction between *X*_1_ and *X*_2_.

**(3) Simulation 3 (“Partially non-linear”):** Similar to Simulation 2 except *f*_0_(**X**) = 10 sin(*πX*_1_*X*_2_)+ 20(*X*_3_ − 0.5)^2^ + 10*X*_4_ + 5*X*_5_ + **X*β***, where the first five elements of ***β*** are zero and only 10% of the remaining linear coefficients are non-zero, drawn randomly from a log-normal distribution similar to Simulation 1.

We compared BayesCOOP with state-of-the-art early, intermediate, and late fusion methods (Ding et al., 2022) in terms of prediction and estimation accuracy, measured by mean squared prediction error (MSPE) and mean squared error (MSE), respectively. For BayesCOOP, we implemented the sampler described in **Algorithm 1** and used the posterior median as our point estimator. To determine the required number of bootstrap pseudo-posterior samples for the coefficients, we gradually increased the number of resamples until the estimates stabilized. For each bootstrap size, we computed the MAP estimate of the coefficients across resamples and monitored the mean, standard deviation, and 2.5% and 97.5% quantiles of each *β* as more bootstraps were added. We continued adding resamples until the Monte Carlo standard errors—defined as the standard deviation divided by the square root of the number of bootstraps—were sufficiently small (*<* 2% of the posterior standard deviation for the mean and *<* 5% of the interquartile range for the quantiles). We also verified that coefficient estimates and confidence interval limits remained stable with additional bootstraps. Once both numerical precision and stability criteria were satisfied, the number of bootstraps was fixed for the final analysis, with 1,100 iterations and the first 100 discarded as burn-in. This configuration provided satisfactory convergence. Hyperparameters were set to their default values (*s*_0_ = 0.05, *s*_1_ = 1), and sensitivity analyses on real data confirmed robustness to these choices (**Supplementary Materials**). For Cooperative Learning, we used the <monospace>R</monospace> package *multiview* (Ding et al., 2022) with default settings and varied the agreement parameter (0, 0.5, 1), corresponding to early, intermediate, and late fusion, respectively.

The simulation results clearly demonstrate that BayesCOOP vastly outperforms published methods in both MSE and MSPE (**Figs. 2-3**) across a range of simulation parameters. Regardless of the sparsity level in the true regression coefficients or the noise level, BayesCOOP achieves either comparable or superior MSE performance relative to early, late, and intermediate fusion. In Simulation 3, which considers a relatively dense setting, BayesCOOP’s performance is comparable to that of Cooperative Learning. However, in sparse settings—regardless of the true association structure—BayesCOOP achieves up to 20 times better out-of-sample prediction accuracy than its competitors, a finding consistent across varying levels of coefficient sparsity and effect sizes. Overall, BayesCOOP provides significant improvements in multiview feature selection and out-of-sample prediction by accommodating the inherent grouping structure within a Bayesian hierarchical framework and effectively propagating uncertainty using pseudo-posteriors.

**Figure 2.**
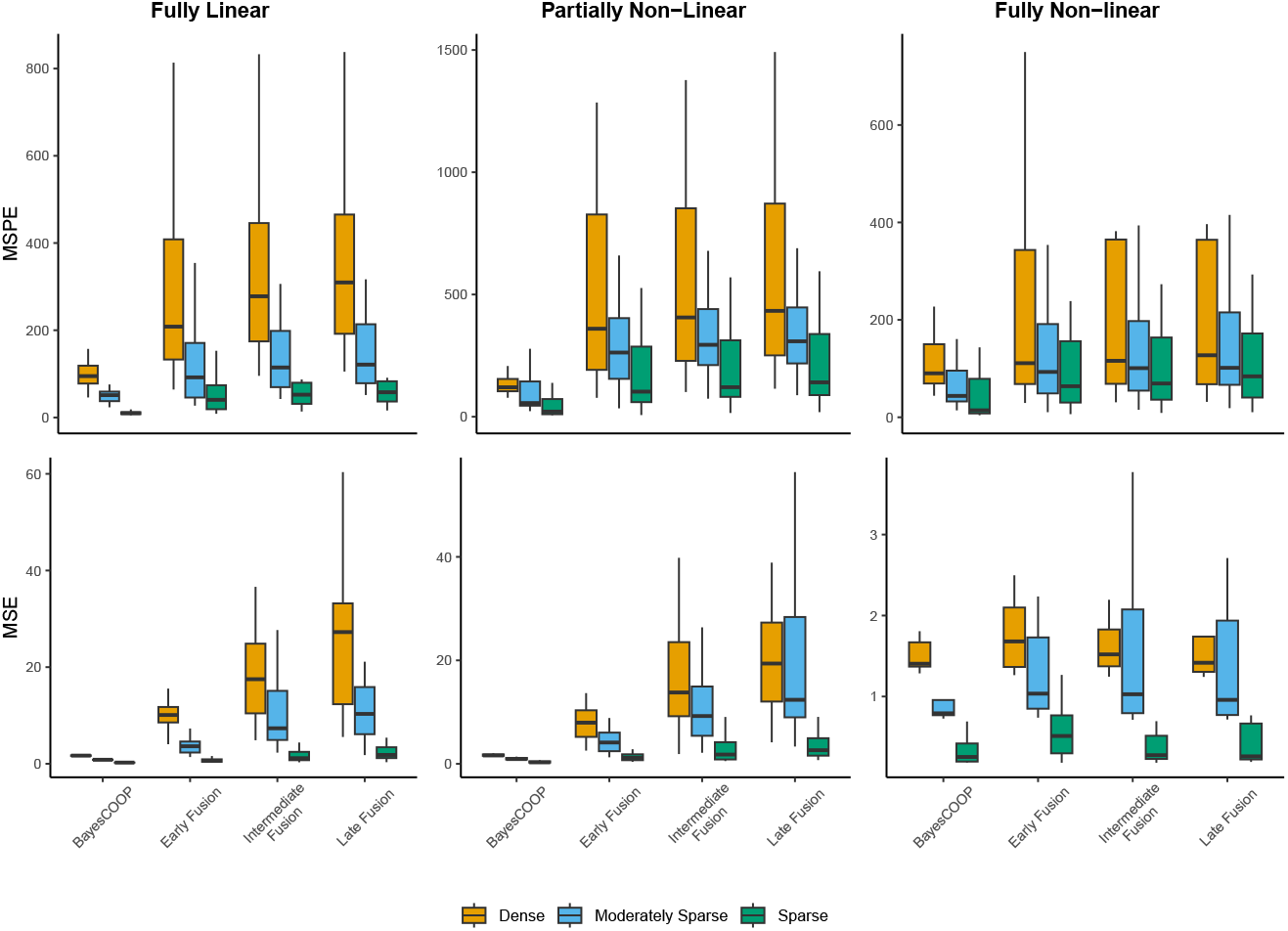
BayesCOOP significantly outperformed published methods in estimation (MSE) and prediction (MSPE) in low SNR (SNR = 5) settings with sample size 500. MSPE values are based on completely held-out test samples, summarized over 25 iterations.

**Figure 3.**
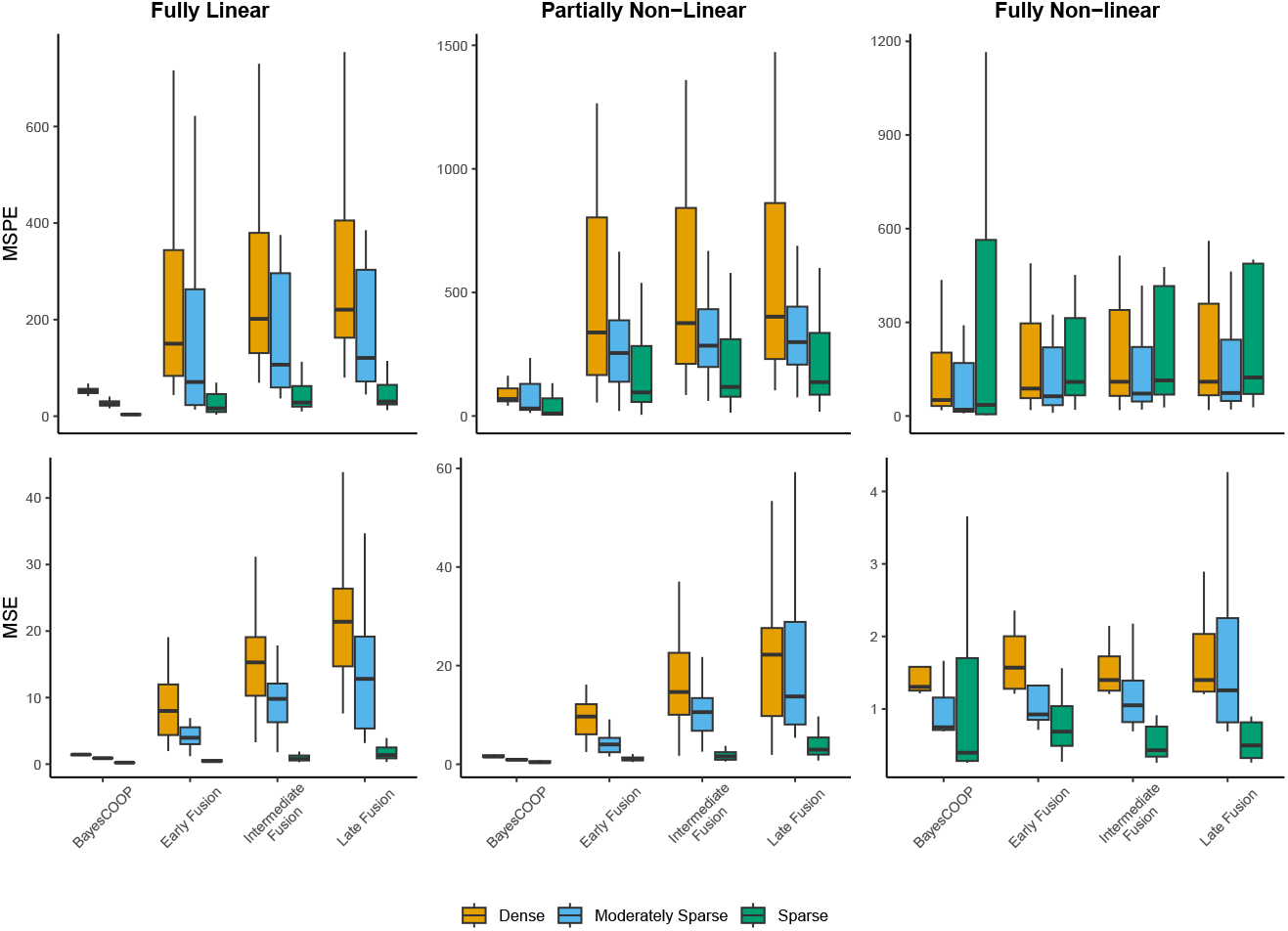
BayesCOOP significantly outperformed published methods in estimation (MSE) and prediction (MSPE) in high SNR (SNR = 10) settings with sample size 500. MSPE values are based on completely held-out test samples, summarized over 25 iterations.

## 4. Real Data Analyses

Following the proof-of-concept validation in synthetic benchmarking, we aimed to examine whether BayesCOOP could facilitate novel discoveries in published studies. To this end, we considered two key questions: (1) Can our proposed method, BayesCOOP, recapitulate published findings? (2) Can BayesCOOP uncover entirely new discoveries through resampling-based Bayesian hierarchical modeling?

To address these questions, we focused on a longitudinal pregnancy dataset from a recent Stanford University study (Stelzer et al., 2021) that profiled proteomic and immunologic adaptations during late gestation. In this cohort, 53 pregnant women receiving routine antepartum care at Lucile Packard Children’s Hospital (Stanford, CA, USA) were followed over the last 100 days of pregnancy. Serial peripheral venous blood samples were collected and profiled using two omics layers—mass cytometry (CyTOF) and proteomics—yielding 2,757 combined features per sample. We analyzed two related outcomes: (1) time to spontaneous labor (**StelzerDOS**), defined as the day of admission for spontaneous labor (contractions occurring at least every 5 minutes, lasting *>* 1 minute, and associated with cervical changes), and (2) gestational age (**StelzerEGA**), determined by the best obstetrical estimate following the American College of Obstetricians and Gynecologists guidelines (Ghaemi et al., 2019). Both datasets included an independent validation set of eight participants, providing an opportunity to rigorously evaluate the generalizability of BayesCOOP.

In all analyses, we first performed independent filtering by removing features with no variability across samples within each omics layer. In addition to BayesCOOP, we applied Cooperative Learning with the agreement parameter fixed at 0.5, the recommended default in the R package *multiview* (Ding et al., 2022). In contrast, BayesCOOP does not require manual tuning of this parameter; instead, the agreement parameter is naturally estimated from the pseudo-posterior samples. As before, we run the corresponding sampler for 1, 000 iterations, discarding the first 100 iterations as burn-in, with the hyperparameters set to their default values: *s*_0_ = 0.05 and *s*_1_ = 1. We report the posterior median as the point estimate for both *ρ* and the regression coefficients ***β***. We additionally applied JAFAR, a recent MCMC-based factor model designed for joint analysis of multiview data (Anceschi et al., 2024). While the application of JAFAR was feasible in the real data analysis, it was not included in the simulation studies due to its high computational cost.

Several key observations are noteworthy. On the pregnancy datasets, BayesCOOP achieved the lowest out-of-sample MSPE on the validation set, significantly outperforming both Cooperative Learning and JAFAR (**Table 1**). Notably, BayesCOOP attained this superior predictive accuracy with a substantially smaller number of selected features—18 vs. 29 in the StelzerDOS dataset and 3 vs. 36 in the StelzerEGA dataset—demonstrating strong generalization through parsimonious modeling. JAFAR was excluded from the feature selection comparison, as it lacks an automatic mechanism for variable selection without post-processing. In the StelzerDOS dataset, BayesCOOP identified four unique features, while showing moderate overlap with Cooperative Learning. The shared features likely represent robust, reproducible biological signals, whereas BayesCOOP’s unique discoveries may reflect complementary associations captured through its Bayesian shrinkage priors.

**Table 1.** Comparison of mean squared prediction error (MSPE) for BayesCOOP, JAFAR (Anceschi et al., 2024), and Cooperative Learning (Ding et al., 2022) in real data analyses: StelzerDOS and StelzerEGA (Stelzer et al., 2021). We also report the number of selected features for BayesCOOP and Cooperative Learning. JAFAR was excluded from the feature selection comparison, as it lacks an automatic mechanism for variable selection without post-processing.

| Dataset | Method | MSPE | Number of Selected Features |
| --- | --- | --- | --- |
| StelzerDOS | BayesCOOP | **493.06** | **18** |
|  | JAFAR | 662.39 | - |
|  | Cooperative Learning | 806.39 | 29 |
| StelzerEGA | BayesCOOP | **6.98** | **3** |
|  | JAFAR | 8.33 | - |
|  | Cooperative Learning | 9.43 | 36 |

While assessing the biological relevance of shared features identified by BayesCOOP and Cooperative Learning in the StelzerDOS dataset, we found that these concordant signals capture proteins with well-characterized patterns around labor onset, reflecting endocrine and stress-response pathways in normal pregnancy (**Fig. 4**). For instance, Activin A and HSP70 both showed higher levels at labor onset, consistent with prior reports linking them to placental differentiation, endocrine activation, and cellular stress adaptation (Barrero et al., 2023; Jee et al., 2021; Tamási et al., 2010). Among the features uniquely identified by BayesCOOP in StelzerDOS, GDF2 (BMP9), FCAR (CD89), ADAM9, and Glucagon represent biologically coherent markers linking immune, vascular, and metabolic adaptation around the transition to labor. GDF2 (BMP9), a vascular-stabilizing ligand of the TGF-*β* superfamily, showed lower levels at labor onset, consistent with its role in early placental endothelial activity (Upton et al., 2020; Aukema et al., 2020). FCAR, which encodes the Fc*α* receptor for IgA, exhibited higher levels at labor onset, consistent with enrichment of CD89^+^ myeloid subsets at the term maternal–fetal interface (Costa et al., 2017). Similarly, ADAM9, a membrane-anchored metalloprotease involved in trophoblast–endothelial interaction and matrix remodeling, showed elevated levels at labor onset, supported by reports of reduced ADAM9 mRNA in preterm placentas and its established role in trophoblast remodeling (Wang et al., 2023; Liu et al., 2024). In contrast, Glucagon, a peptide hormone regulating glucose metabolism, showed reduced levels at labor onset, consistent with tighter endocrine control and metabolic transition toward efficient energy utilization near delivery (Leblanc et al., 1976; Perlman et al., 1988).

**Figure 4.**
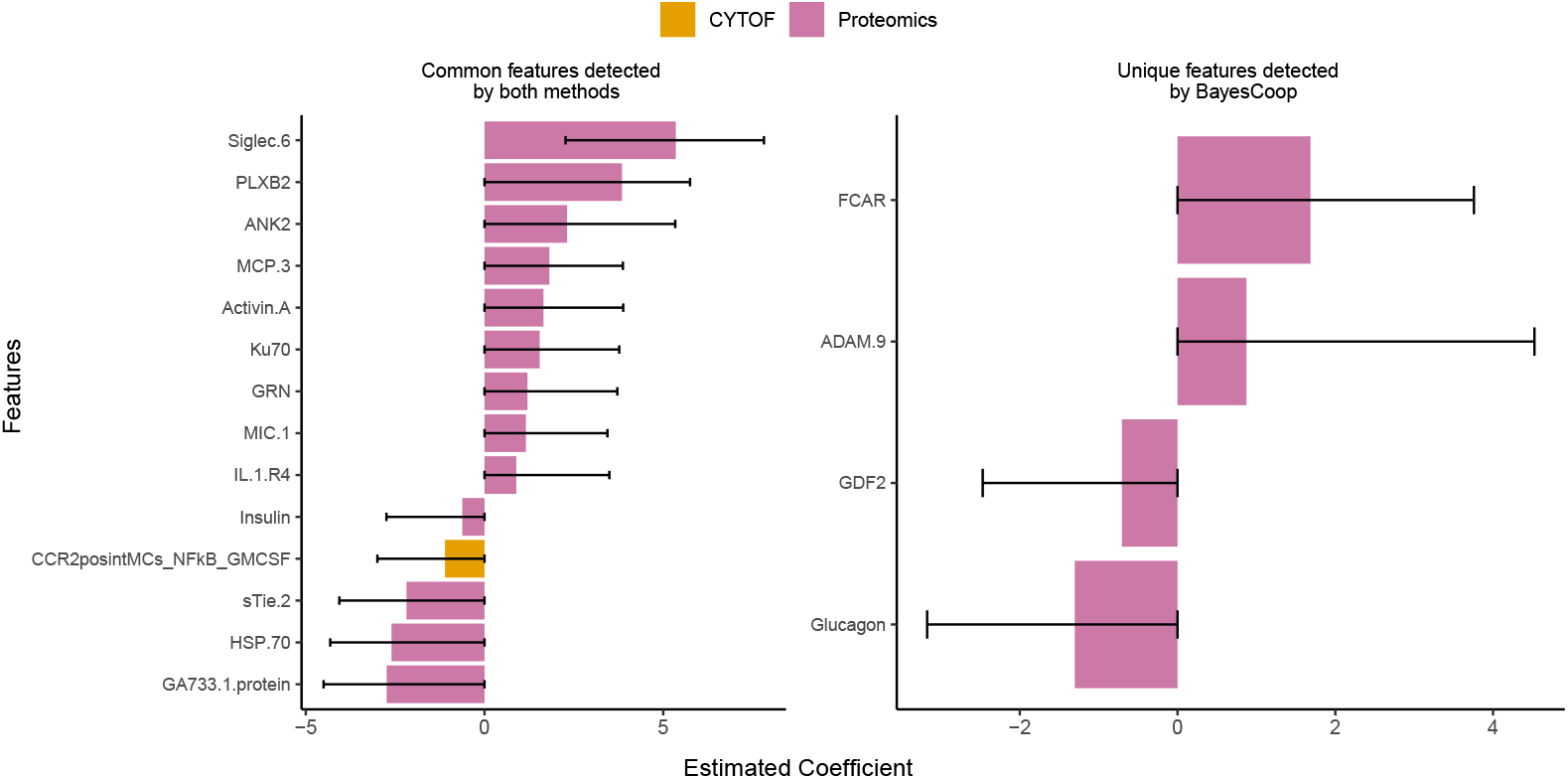
BayesCOOP identifies unique features not detected by published methods in the StelzerDOS dataset, including several biologically relevant candidates worthy of follow-up experiments. Effect sizes (posterior medians and credible intervals) for selected multiview features uniquely identified by BayesCOOP (right) and those detected by both Cooperative Learning (Ding et al., 2022) and BayesCOOP (left) are shown.

In the StelzerEGA dataset, BayesCOOP identified three proteomic features: MIC-1 (GDF15), Siglec-6, and PLXB2. Interestingly, all three markers overlapped with those detected by Cooperative Learning and were also present in the StelzerDOS analysis, indicating strong reproducibility across pregnancy outcomes (**Fig. S3**). MIC-1 (GDF15) showed a positive association with gestational age, consistent with its progressive rise throughout pregnancy and its established role in placental endocrine signaling and metabolic adaptation preceding delivery (Høgh et al., 2025). Siglec-6 also increased with gestational age, consistent with its upregulation in term trophoblasts and higher expression in placentas at later gestation, implicating it in cytokine- and leptin-mediated signaling at the maternal–fetal interface (Brinkman-Van der Linden et al., 2007). PLXB2 similarly trended positively with gestational age, in line with its role in trophoblast migration, vascular morphogenesis, and maintenance of epithelial integrity in the endometrium, where it localizes to apical and lateral membranes to support barrier cohesion (Singh and Aplin, 2015). Although these proteomic signatures marking the transition from mid-to late gestation remain exploratory and warrant validation in larger, longitudinal pregnancy cohorts, their concordance with established biological mechanisms underscores BayesCOOP’s ability to extract mechanistically meaningful patterns from high-dimensional multimodal data.

As a final evaluation using real data, we examined the posterior predictive intervals generated by BayesCOOP in both the training and validation cohorts (**Figs. 5;S4**). The predicted values closely aligned with the observed outcomes, while the widths of the posterior intervals appropriately reflected varying levels of predictive uncertainty across subjects. In the training cohort, the posterior intervals were generally narrower, consistent with stronger model fit and lower residual variability. In contrast, the validation cohort exhibited slightly wider intervals, reflecting greater uncertainty when extrapolating to unseen data. This comparison highlights that BayesCOOP remains well-calibrated across cohorts, effectively adapting interval width to data uncertainty. Overall, this analysis underscores the importance of explicitly quantifying uncertainty—particularly in the presence of substantial population heterogeneity—since relying solely on point estimates can lead to misleading or overconfident conclusions.

**Figure 5.**
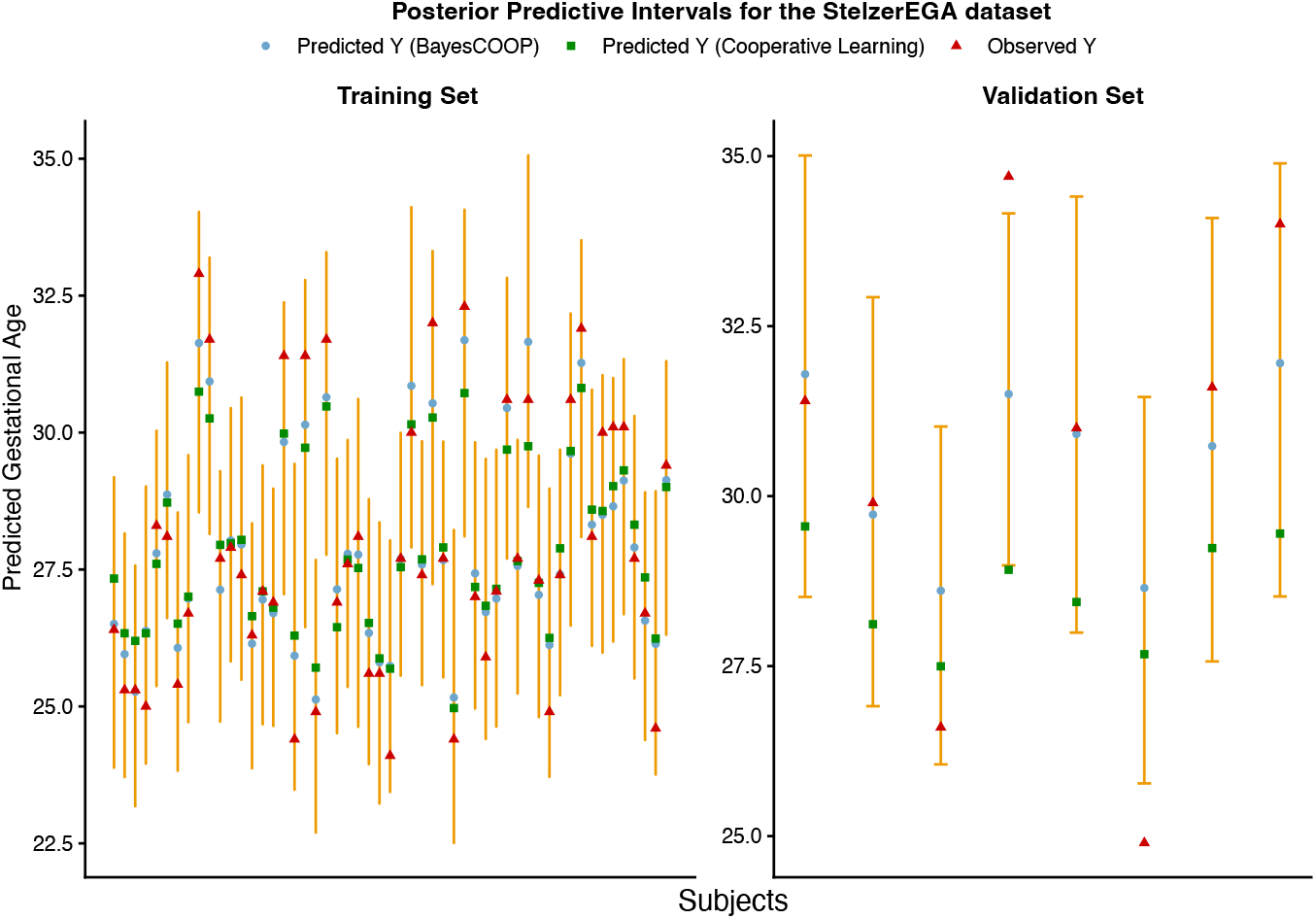
BayesCOOP delivers well-calibrated and more accurate predictions of gestational age in both the training and validation sets of the StelzerEGA dataset (Stelzer et al., 2021), outperforming Cooperative Learning (Ding et al., 2022), which provides no quantification of uncertainty in its estimates.

## 5. Discussion

BayesCOOP provides the first scalable Bayesian framework for multimodal integration, unifying group-aware sparsity with principled uncertainty quantification. By leveraging pseudo-posterior approximation within a Bayesian bootstrap formulation, BayesCOOP embeds a fast, deterministic algorithm that delivers stable posterior summaries without the computational burden of MCMC. Theoretical guarantees ensure posterior consistency, while empirical studies demonstrate substantial gains in estimation and prediction accuracy over published methods. As part of our ongoing research, we are extending BayesCOOP to accommodate spatial multimodal analysis (Chen et al., 2025), multi-study integration (Liu and Zhong, 2025), transfer learning (Tian and Feng, 2023), pre-training (Craig et al., 2025), and causal multimodal inference (Jiang et al., 2025), thereby broadening its methodological scope and translational impact. Taken together, these developments position BayesCOOP as a unifying and extensible foundation for scalable, uncertainty-aware learning across complex multimodal domains.

## Supporting Information

The proofs of Theorems 1 and 2 referenced in Section 2.3, and additional tables and figures referenced in Sections 3 and 4 are available in the **Supplementary Materials**. The BayesCOOP R package is available at https://github.com/himelmallick/BayesCOOP.

## References

Anceschi, N., Ferrari, F., Dunson, D. B., and Mallick, H. (2024). Bayesian joint additive factor models for multiview learning. arXiv preprint 2406.00778.

Aukema, S. M., Ten Brinke, G. A., Timens, W., Vos, Y. J., Accord, R. E., Kraft, K. E., Santing, M. J., Morssink, L. P., Streefland, E., van Diemen, C. C., et al. (2020). A homozygous variant in growth and differentiation factor 2 (gdf2) may cause lymphatic dysplasia with hydrothorax and nonimmune hydrops fetalis. American Journal of Medical Genetics Part A 182, 2152–2160.

Barrero, J. A., Villamil-Camargo, L. M., Imaz, J. N., Arciniegas-Villa, K., and Rubio-Romero, J. A. (2023). Maternal serum activin a, inhibin a and follistatin-related proteins across preeclampsia: insights into their role in pathogenesis and prediction. Journal of mother and child 27, 119.

Brinkman-Van der Linden, E. C., Hurtado-Ziola, N., Hayakawa, T., Wiggleton, L., Benirschke, K., Varki, A., and Varki, N. (2007). Human-specific expression of siglec-6 in the placenta. Glycobiology 17, 922–931.

Chalise, P., Raghavan, R., and Fridley, B. L. (2016). Intersim: Simulation tool for multiple integrative ‘omic datasets’. Computer methods and programs in biomedicine 128, 69–74.

Chen, J. G., Chávez-Fuentes, J. C., O’Brien, M., Xu, J., Ruiz, E. C., Wang, W., Amin, I., Sheridan, J. P., Shin, S. C., Hasyagar, S. V., et al. (2025). Giotto suite: a multiscale and technology-agnostic spatial multiomics analysis ecosystem. Nature Methods pages 1–13.

Chung, R.-H. and Kang, C.-Y. (2019). A multi-omics data simulator for complex disease studies and its application to evaluate multi-omics data analysis methods for disease classification. GigaScience 8, giz045.

Costa, M. L., Robinette, M. L., Bugatti, M., Longtine, M. S., Colvin, B. N., Lantelme, E., Vermi, W., Colonna, M., Nelson, D. M., and Cella, M. (2017). Two distinct myeloid subsets at the term human fetal–maternal interface. Frontiers in immunology 8, 1357.

Craig, E., Pilanci, M., Le Menestrel, T., Narasimhan, B., Rivas, M. A., Gullaksen, S.-E., Dehghannasiri, R., Salzman, J., Taylor, J., and Tibshirani, R. (2025). Pretraining and the lasso. Journal of the Royal Statistical Society Series B: Statistical Methodology page qkaf050.

De Vito, R., Bellio, R., Trippa, L., and Parmigiani, G. (2021). Bayesian multistudy factor analysis for high-throughput biological data. The annals of applied statistics 15, 1723– 1741.

Ding, D. Y., Li, S., Narasimhan, B., and Tibshirani, R. (2022). Cooperative learning for multiview analysis. Proceedings of the National Academy of Sciences 119, e2202113119.

Friedman, J. H. (1991). Multivariate adaptive regression splines. The annals of statistics 19, 1–67.

George, E. I. and McCulloch, R. E. (1993). Variable selection via gibbs sampling. Journal of the American Statistical Association 88, 881–889.

Ghaemi, M. S. et al. (2019). Multiomics modeling of the immunome, transcriptome, microbiome, proteome and metabolome adaptations during human pregnancy. Bioinformatics 35, 95–103.

Grabski, I. N., De Vito, R., Trippa, L., and Parmigiani, G. (2023). Bayesian combinatorial multistudy factor analysis. The annals of applied statistics 17, 2212.

Hansen, B., Avalos-Pacheco, A., Russo, M., and De Vito, R. (2022). A variational bayes approach to factor analysis. In International Conference on Bayesian Statistics in Action, pages 15–21. Springer.

Hansen, B., Avalos-Pacheco, A., Russo, M., and De Vito, R. (2025). Fast variational inference for bayesian factor analysis in single and multi-study settings. Journal of Computational and Graphical Statistics 34, 96–108.

Høgh, S., Borgsted, C., Hegaard, H. K., Renault, K. M., Ekelund, K., Bruzzone, S. E., Clemmensen, C., Klein, A. B., and Frokjaer, V. G. (2025). Growth differentiation factor 15 during pregnancy and postpartum as captured in blood, cerebrospinal fluid and placenta: A cohort study on associations with maternal mental health. Psychoneuroendocrinology 171, 107212.

Huggins, J. H. and Miller, J. W. (2024). Reproducible parameter inference using bagged posteriors. Electronic Journal of Statistics 18, 1549–1585.

Jee, B., Dhar, R., Singh, S., and Karmakar, S. (2021). Heat shock proteins and their role in pregnancy: Redefining the function of “old rum in a new bottle”. Frontiers in Cell and Developmental Biology 9, 648463.

Jiang, H., Miao, X., Thairu, M. W., Beebe, M., Grupe, D. W., Davidson, R. J., Handelsman, J., and Sankaran, K. (2025). Multimedia: multimodal mediation analysis of microbiome data. Microbiology Spectrum 13, e01131–24.

Leblanc, H., Anderson, J., and Yen, S. (1976). Glucagon secretion in late pregnancy and the puerperium. American Journal of Obstetrics and Gynecology 125, 708–710.

Liu, J., Wang, Y., Zhang, S., Sun, L., and Shi, Y. (2024). Adam9 deubiquitination induced by usp22 suppresses proliferation, migration, invasion, and epithelial-mesenchymal transition of trophoblast cells in preeclampsia. Placenta 146, 50–57.

Liu, W. and Zhong, Q. (2024). High-dimensional covariate-augmented overdispersed poisson factor model. Biometrics 80, ujae031.

Liu, W. and Zhong, Q. (2025). High-dimensional multi-study multi-modality covariate-augmented generalized factor model. Biometrics 81, ujaf107.

Mallick, H., Porwal, A., Saha, S., Basak, P., Svetnik, V., and Paul, E. (2024). An integrated bayesian framework for multi-omics prediction and classification. Statistics in Medicine 43, 983–1002.

Mauri, L. and Dunson, D. B. (2025). Inference on covariance structure in high-dimensional multi-view data. arXiv e-prints pages arXiv–2509.

Monzó, C., Martínez-Mira, C., Arzalluz-Luque, Á., Conesa, A., and Tarazona, S. (2018). Mosim: bulk and single-cell multi-layer regulatory network simulator.

Nie, L. and Ročková, V. (2023). Bayesian bootstrap spike-and-slab lasso. Journal of the American Statistical Association 118, 2013–2028.

Perlman, R., Halabi, C., Bick, T., and Hochberg, Z. (1988). The human placenta as a target tissue for glucagon. Biochemical and biophysical research communications 151, 1019–1024.

Ročková, V. and George, E. I. (2018). The spike-and-slab lasso. Journal of the American Statistical Association 113, 431–444.

Rubin, D. B. (1981). The bayesian bootstrap. The annals of statistics pages 130–134.

Samorodnitsky, S., Wendt, C. H., and Lock, E. F. (2024). Bayesian simultaneous factorization and prediction using multi-omic data. Computational Statistics & Data Analysis 197, 107974.

Singh, H. and Aplin, J. (2015). Endometrial apical glycoproteomic analysis reveals roles for cadherin 6, desmoglein-2 and plexin b2 in epithelial integrity. Molecular human reproduction 21, 81–94.

Srivastava, S., Li, C., and Dunson, D. B. (2018). Scalable bayes via barycenter in wasserstein space. Journal of Machine Learning Research 19, 1–35.

Stelzer, I. A. et al. (2021). Integrated trajectories of the maternal metabolome, proteome, and immunome predict labor onset. Science Translational Medicine 13,.

Tamási, L., Bohács, A., Tamási, V., Stenczer, B., Prohászka, Z., Rigó Jr, J., Losonczy, G., and Molvarec, A. (2010). Increased circulating heat shock protein 70 levels in pregnant asthmatics. Cell Stress and Chaperones 15, 295–300.

Tang, Z., Shen, Y., Li, Y., Zhang, X., Wen, J., Qian, C., Zhuang, W., Shi, X., and Yi, N. (2018). Group spike-and-slab lasso generalized linear models for disease prediction and associated genes detection by incorporating pathway information. Bioinformatics 34, 901–910.

Tian, Y. and Feng, Y. (2023). Transfer learning under high-dimensional generalized linear models. Journal of the American Statistical Association 118, 2684–2697.

Upton, P. D., Park, J. E., De Souza, P. M., Davies, R. J., Griffiths, M. J., Wort, S. J., and Morrell, N. W. (2020). Endothelial protective factors bmp9 and bmp10 inhibit ccl2 release by human vascular endothelial cells. Journal of cell science 133, jcs239715.

Wang, X., Liu, A., Hou, D., Dong, X., Chu, C., Ju, W., Zhang, J., Jia, Y., Yang, X., Ji, Y., et al. (2023). Epigenetic impact of long non-coding rna lnc-adam9 on extracellular matrix pathway in preterm syndrome through down-regulation of mrna-adam9. Journal of Translational Genetics and Genomics 7, 126–140.

Xu, Y. and McCord, R. P. (2022). Diagonal integration of multimodal single-cell data: potential pitfalls and paths forward. Nature Communications 13, 3505.

Zappia, L., Phipson, B., and Oshlack, A. (2017). Splatter: simulation of single-cell rna sequencing data. Genome biology 18, 174.

