## Supplementary Materials for "Bayesian Cooperative Learning for Multimodal Integration"

#### A BayesCOOP for Generalized Outcomes

In this section, motivated by the ubiquity of binary, count, and survival outcomes in multiview studies [Ding et al., 2022; Mallick et al., 2024], we extend BayesCOOP to accommodate generalized outcomes from the exponential family (e.g., binary and count data) as well as parametric survival models such as the accelerated failure time (AFT) model. This extension enables flexible and coherent modeling of diverse response types while preserving the cooperative structure across multiple data views. As before, we consider a setup involving $d$ views, represented as $\boldsymbol{X}^{(1)}, \ldots, \boldsymbol{X}^{(d)}$ with $\boldsymbol{X}^{(i)} \in \mathbb{R}^{n \times p_i}, \forall i = 1, \ldots, d$ and an outcome variable $\boldsymbol{y} = (y_1, \ldots, y_n)^\top$. Also define, $\boldsymbol{X} = \left[\boldsymbol{X}^{(1)}, \ldots, \boldsymbol{X}^{(d)}\right]$ where $\boldsymbol{X}_j^{(i)} \in \mathbb{R}^{p_i} \quad \forall j = 1, \ldots, n, \quad \text{and} \quad i = 1, \ldots, d.$

#### A.1 Probit Models for Binary Outcomes

Consider a binary response variable $\boldsymbol{y} = (y_1, \ldots, y_n)^\top$, where each $y_i \in \{0, 1\}$. We assume

$$(y_i \mid \pi_i) \overset{\text{ind}}{\sim} \text{Bernoulli}(\pi_i), \qquad \pi_i = \Phi(\eta_i), \qquad \eta_i = \sum_{m=1}^{d} \boldsymbol{X}_i^{(m)\top} \boldsymbol{\beta}_m,$$

where $\Phi(\cdot)$ denotes the cumulative distribution function of a standard normal distribution, and $\boldsymbol{\beta}_m \in \mathbb{R}^{p_m}$ is the coefficient vector corresponding to the $m$-th view.

Following the classical data augmentation approach of Albert and Chib [1993], we introduce latent Gaussian variables $\boldsymbol{z} = (z_1, \ldots, z_n)^\top$ such that

$$z_i \overset{\text{ind}}{\sim} \mathcal{N}\left(\sum_{m=1}^{d} \boldsymbol{X}_i^{(m)\top} \boldsymbol{\beta}_m, \sigma^2\right), \qquad y_i = \begin{cases} 1, & \text{if } z_i > 0, \\ 0, & \text{if } z_i \leq 0. \end{cases}$$

This latent variable representation converts the binary likelihood into a conditionally Gaussian form, which simplifies both parameter estimation and Bayesian computation.

Conditional on $\boldsymbol{z}$, we define the *regularized multiview likelihood* as:

$$\mathcal{L}(\boldsymbol{\beta} \mid \boldsymbol{z}, \boldsymbol{y}, \boldsymbol{X}^{(1)}, \ldots, \boldsymbol{X}^{(d)}, \rho, \sigma^2) \propto \exp\left[-\frac{1}{2\sigma^2}\left\{\|\boldsymbol{z} - \sum_{m=1}^{d} \boldsymbol{X}^{(m)} \boldsymbol{\beta}_m\|_2^2 + \rho \sum_{m<\ell} \|\boldsymbol{X}^{(m)} \boldsymbol{\beta}_m - \boldsymbol{X}^{(\ell)} \boldsymbol{\beta}_\ell\|_2^2\right\}\right], \tag{A.1.1}$$

where $\rho > 0$ serves as an *agreement parameter* that regulates information sharing across views by penalizing discrepancies between view-specific predictions [Ding et al., 2022]. Rewriting

$$\widetilde{\boldsymbol{z}} = (\boldsymbol{z}^\top, \boldsymbol{0}^\top, \ldots, \boldsymbol{0}^\top)^\top \in \mathbb{R}^{ns},$$

with $s = 1 + \frac{d(d-1)}{2}$, and an augmented design matrix $\widetilde{\boldsymbol{X}}$, similar to the one in the main paper, the likelihood in (A.1.1) can be equivalently rewritten as a standard Gaussian likelihood on the augmented data:

$$\mathcal{L}(\boldsymbol{\beta} \mid \widetilde{\boldsymbol{z}}, \boldsymbol{y}, \widetilde{\boldsymbol{X}}, \rho, \sigma^2) \propto \exp\left(-\frac{1}{2\sigma^2}\|\widetilde{\boldsymbol{z}} - \widetilde{\boldsymbol{X}}\boldsymbol{\beta}\|_2^2\right).$$

Given this augmentation scheme and prior specification as in equation 5 in the manuscript, **Algorithm 1** can be applied directly, with an additional step to update the latent variables $\boldsymbol{z}$ from their posterior conditional distributions:

$$z_i \mid \boldsymbol{\beta}, \boldsymbol{y}, \boldsymbol{X}^{(1)}, \ldots, \boldsymbol{X}^{(d)}, \rho, \sigma^2 \overset{\text{ind}}{\sim} \begin{cases} \mathcal{N}\left(\sum_{m=1}^{d} \boldsymbol{X}_i^{(m)\top} \boldsymbol{\beta}_m, \sigma^2\right) \text{ truncated below at 0,} & \text{if } y_i = 1, \\ \mathcal{N}\left(\sum_{m=1}^{d} \boldsymbol{X}_i^{(m)\top} \boldsymbol{\beta}_m, \sigma^2\right) \text{ truncated above at 0,} & \text{if } y_i = 0. \end{cases}$$

This latent variable sampling step is carried out at each iteration of the Bayesian bootstrap procedure in **Algorithm 1**, allowing the model to capture both posterior uncertainty and cross-view dependencies for binary outcomes in a coherent and computationally tractable manner.

##### A.2 Accelerated Failure Time Models for Survival Outcomes

Consider a typical survival analysis setting with the outcome variable $\boldsymbol{y} = (\boldsymbol{t}, \boldsymbol{c})$ where $\boldsymbol{t} = (t_1, \ldots, t_n)^\top \in \mathbb{R}^n$ and $\boldsymbol{c} = (c_1, \ldots, c_n)^\top \in \mathbb{R}^n$ with $t_i$ and $c_i$ denoting the survival time and the censoring time of the $i$-th subject $(i = 1, \ldots, n)$ respectively. We model the logarithm of the survival time using the accelerated failure time (AFT) model:

$$\log(t_i) = \sum_{m=1}^{d} \boldsymbol{X}_i^{(m)\top} \boldsymbol{\beta}_m + \epsilon_i, \quad i = 1, \ldots, n,$$

where $\boldsymbol{\epsilon} = (\epsilon_1, \ldots, \epsilon_n)^\top$ is the vector of error terms. Assuming $\epsilon_i \sim N(0, \sigma^2)$ leads to the log-normal AFT model, although other distributions such as the $t$-distribution may also be used for robustness [Kleinbaum and Klein, 2006].

Suppose the outcomes are subjected to right censoring. Therefore, the observed survival time is given by $t_i^* = \min(t_i, c_i)$, and the corresponding censoring indicator is $u_i = \mathbb{I}\{t_i \leq c_i\}$. Following Sha et al., 2006 and Maity et al., 2019, we adopt a similar data augmentation strategy to impute the latent log-survival times. Specifically, we define the augmented variable $\boldsymbol{w} = (w_1, \ldots, w_n)^\top$ with $w_i = \log(t_i)$ such that,

$$w_i = \begin{cases} \log(t_i^*), & \text{if } u_i = 1 \text{ (event observed)}, \\ > \log(t_i^*), & \text{if } u_i = 0 \text{ (right censored)}, \end{cases}$$

Here, $w_i$ coincides with the observed log-survival time for uncensored subjects and is constrained to exceed $\log(t_i^*)$ when the event is censored. This augmented variable $w_i$ serves as the latent log-survival time in the AFT model, enabling standard regression-based inference and posterior sampling. Now, the *regularized multiview likelihood function*, which introduces an agreement parameter that penalizes discrepancies between predicted responses across pairs of views [Ding et al., 2022] is given by,

$$\mathcal{L}(\boldsymbol{\beta} \mid \boldsymbol{w}, \widetilde{\boldsymbol{X}}, \rho, \sigma^2) \propto \exp\left[ -\frac{1}{2\sigma^2} \left\{ \left\| \boldsymbol{w} - \sum_{m=1}^{d} \boldsymbol{X}^{(m)}\boldsymbol{\beta}_m \right\|_2^2 + \rho \sum_{m<\ell} \left\| \boldsymbol{X}^{(m)}\boldsymbol{\beta}_m - \boldsymbol{X}^{(\ell)}\boldsymbol{\beta}_\ell \right\|_2^2 \right\} \right]. \quad \text{(A.2.1)}$$

We define the augmented response vector as $\widetilde{\boldsymbol{w}} = (\boldsymbol{w}^\top, \mathbf{0}^\top, \ldots, \mathbf{0}^\top)^\top \in \mathbb{R}^{ns}$, where $s = 1 + \frac{d(d-1)}{2}$. Let $\widetilde{\boldsymbol{X}}$ denote the corresponding augmented design matrix constructed analogously to that in the main paper. Under this formulation, the likelihood in (A.2.1) can be expressed as a standard Gaussian likelihood based on the augmented data:

$$\mathcal{L}(\boldsymbol{\beta} \mid \widetilde{\boldsymbol{w}}, \widetilde{\boldsymbol{X}}, \rho, \sigma^2) \propto \exp\left( -\frac{1}{2\sigma^2} \left\| \widetilde{\boldsymbol{w}} - \widetilde{\boldsymbol{X}}\boldsymbol{\beta} \right\|_2^2 \right).$$

Given this augmentation scheme and the prior specification in equation 5 of the manuscript, Algorithm 1 can be implemented directly, with an additional step to update the latent variables $w_i$ by drawing them from their full conditional distributions $\mathcal{N}\left( \sum_{m=1}^{d} \boldsymbol{X}_i^{(m)\top} \boldsymbol{\beta}_m, \sigma^2 \right)$ left truncated at $\log(t_i^*)$ for each iteration of the Bayesian bootstrap procedure.

#### A.3 Least Square Approximation for Generalized Outcomes

Finally, within the realm of reasonable large-sample approximation, similar algorithms can be used to fit BayesCOOP for generalized outcomes from the exponetial family, extending the BayesCOOP to more complex models such as count models including zero-inflated models, and generalized linear models (GLMs). Let us denote by $L(\boldsymbol{\beta})$ the negative log-likelihood. Following Wang and Leng [2007], $L(\boldsymbol{\beta})$ can be approximated by least-squares approximation (LSA) as follows:

$$L(\boldsymbol{\beta}) \approx \frac{1}{2}(\boldsymbol{\beta} - \tilde{\boldsymbol{\beta}})' \hat{\boldsymbol{\Sigma}} (\boldsymbol{\beta} - \tilde{\boldsymbol{\beta}}),$$

where $\tilde{\boldsymbol{\beta}}$ is the MLE of $\boldsymbol{\beta}$ and $\hat{\Sigma}^{-1} = \partial^2 L(\boldsymbol{\beta}) / \partial \boldsymbol{\beta}^2$. Therefore, for a general model, the conditional distribution of $\boldsymbol{y}$ is given by

$$\boldsymbol{y} | \boldsymbol{\beta} \sim \exp\left\{ -\frac{1}{2\sigma^2} (\boldsymbol{\beta} - \tilde{\boldsymbol{\beta}})' \hat{\boldsymbol{\Sigma}} (\boldsymbol{\beta} - \tilde{\boldsymbol{\beta}}) \right\}.$$

This allows the general likelihoods to be similarly represented using the hierarchies introduced in the manuscript [Leng et al., 2014], yielding tractable full conditional distributions.

### B Proof of Theorems

In this section, we outline the theoretical framework for establishing the properties of BayesCOOP by extending the key concepts from Nie and Ročková [2023] to the multiview setting considered in the current article.

#### B.1 Setup

Recall the model $\widetilde{\boldsymbol{y}} = \widetilde{\boldsymbol{X}}\boldsymbol{\beta} + \boldsymbol{\epsilon}$ based on the augmented data comprising $ns$ observations, with $\boldsymbol{\epsilon} \sim N(0, \sigma^2 I_{ns})$, and $p$-dimensional coefficient vector $\boldsymbol{\beta}$ where $p = \sum_{j=1}^{d} p_j$. The weighted log-posterior (upto constants) of $\boldsymbol{\beta}$ is then given by

$$Q(\boldsymbol{\beta}) = -\frac{1}{2\sigma^2}\|\boldsymbol{W}(\widetilde{\boldsymbol{y}} - \widetilde{\boldsymbol{X}}\boldsymbol{\mu}) - \boldsymbol{W}\widetilde{\boldsymbol{X}}\boldsymbol{\beta}\|_2^2 + \sum_{j=1}^{d}\sum_{k=1}^{p_d} \pi(\beta_{jk} \mid \gamma_{jk}, s_0), \tag{B.1.1}$$

where $\pi(\beta_{jk} \mid \gamma_{jk}, s_0)$ denotes the grouped spike-and-slab log-prior of the $k$-th coefficient belonging to the $j$-th group, and we define $\boldsymbol{W} = \text{diag}(\sqrt{w_1}, \ldots, \sqrt{w_{ns}})$. The MAP estimator $\hat{\boldsymbol{\beta}}$ maximizes $Q(\boldsymbol{\beta})$ and the pseudo-posterior draws are given by $\tilde{\boldsymbol{\beta}} = \hat{\boldsymbol{\beta}} + \boldsymbol{\mu}$, where $\boldsymbol{\mu}$ represents the jitter in prior mean of $\boldsymbol{\beta}$. Let $q = \|\boldsymbol{\beta_0}\|$ be the sparsity of the true coefficient $\boldsymbol{\beta_0}$ and $\hat{q} = \|\hat{\boldsymbol{\beta}}\|$ be the number of selected coordinates in the MAP estimate.

#### B.2 Proof of Theorem 1

Since $\hat{\boldsymbol{\beta}}$ maximizes $Q(\boldsymbol{\beta})$, we have $0 \geq Q(\boldsymbol{\beta_0}) - Q(\hat{\boldsymbol{\beta}})$. We decompose this difference into

$$0 \geq \|\boldsymbol{W}\widetilde{\boldsymbol{X}}\boldsymbol{\Theta}\|_2^2 - 2\big(\boldsymbol{W}\boldsymbol{\epsilon} - \boldsymbol{W}\widetilde{\boldsymbol{X}}\boldsymbol{\mu}\big)^\top \boldsymbol{W}\widetilde{\boldsymbol{X}}\boldsymbol{\Theta} + 2\sigma^2 \sum_{j=1}^{d}\sum_{k=1}^{p_j}\big\{\pi(\beta_{0,jk}) - \pi(\hat{\beta}_{jk})\big\}, \tag{B.2.1}$$

where $\boldsymbol{\Theta}$ is defined in Lemma A.6 [Nie and Ročková, 2023]. Let us first consider the second term in (B.2.1). Extending Lemma A.7 of Nie and Ročková [2023], it can be shown that

$$-2\big(\boldsymbol{W}\boldsymbol{\epsilon} - \boldsymbol{W}\widetilde{\boldsymbol{X}}\boldsymbol{\mu}\big)^\top \boldsymbol{W}\widetilde{\boldsymbol{X}}\boldsymbol{\Theta} \geq -2\big(\sqrt{M}\,\eta^*\Delta_U + \sigma^2/s_1\big)\|\boldsymbol{\Theta}\|_1, \tag{B.2.2}$$

where $\Delta_U = \sqrt{2\,ns\,\sigma^2\log(1/p^*(0))} + \sigma^2/s_1$. Furthermore, the last term in (B.2.1) is greater than or equal to $\widehat{q}\,b_w + (\widehat{q} - q)\log[1/p^*(0)]$. Definition A.2 and Lemma A.6 of Nie and Ročková [2023] then yield

$\|\mathbf{\Theta}\|_1 \le 2\|\mathbf{\Theta}\|_2\sqrt{\widehat{q}+q}$ and $\|\boldsymbol{W}\widetilde{\boldsymbol{X}}\mathbf{\Theta}\|_2^2 \ge m\,c^2(\eta^*;\boldsymbol{\beta_0})\,ns\,\|\mathbf{\Theta}\|_2^2$. Combining these bounds, we get

$$(\widehat{q}-q)\log\left[1/p^*(0)\right] + \widehat{q}\,b_w \le \frac{\left(\sqrt{M}\,\eta^*\Delta_U + \sigma^2/s_1\right)^2}{2\,m\,\sigma^2\,c^2(\eta^*;\beta_0)\,ns}\,(\widehat{q}+q) \equiv D(\hat{q}+q), \tag{B.2.3}$$

which together with Lemma A.4 and A.5, and conditions (i)-(v) in Nie and Ročková [2023], yield that, the recovered model-size $\hat{q}$ by BayesCOOP satisfies,

$$\lim_{n\to\infty}\mathbb{E}_{\boldsymbol{\beta}_0}\mathbb{P}_{\boldsymbol{\mu},\boldsymbol{w}}\left(\|\widetilde{\boldsymbol{\beta}}_w^{\boldsymbol{\mu}}-\boldsymbol{\mu}\|_0 \le q(1+K)\,\middle|\,\widetilde{\boldsymbol{y}}\right) = 1,$$

where $K := \frac{2D}{1-D}$.

#### B.3 Proof of Theorem 2

The proof of this theorem follows the same arguments as the proofs of Theorem 8 in Ročková and George [2018] and Theorem 4.3 in Nie and Ročková [2023]. We begin with the triangle inequality:

$$\|\widetilde{\boldsymbol{\beta}}-\boldsymbol{\beta_0}\|_2 \le \|\widetilde{\boldsymbol{\beta}}-\hat{\boldsymbol{\beta}}\|_2 + \|\hat{\boldsymbol{\beta}}-\boldsymbol{\beta_0}\|_2.$$

Thereafter, the proof proceeds generally through two key steps. First, we need to have a high probability bound on the quantity $\|\mathbf{\Theta}\|_2 = \|\hat{\boldsymbol{\beta}}-\boldsymbol{\beta_0}\|_2$. Since $Q(\widehat{\boldsymbol{\beta}}) \ge Q(\boldsymbol{\beta_0})$, and using Lemma A.6 and Lemma A.7 and conditions (i) and (ii) of the proof of Theorem 4.3 in Nie and Ročková [2023], we obtain,

$$0 \ge \|\boldsymbol{W}\widetilde{\boldsymbol{X}}\mathbf{\Theta}\|_2^2 - 2\left(\sqrt{M}\eta^*\Delta^U + \frac{1}{s_1}\right)\|\mathbf{\Theta}\|_1 + 2q\log p^*(0), \tag{B.3.1}$$

where $\eta^*$ and $\Delta^U$ are defined in the statement of the Theorem 2. Now, from the proof of Theorem 1, since $\|\mathbf{\Theta}\|_0 \le (1+K)q$ and $(a-2b)^2 \ge 0$, we can provide a bound for $2\left(\sqrt{M}\eta^*\Delta^U+\lambda_1\right)\|\mathbf{\Theta}\|_1$ as:

$$2\left(\sqrt{M}\eta^*\Delta^U + \frac{1}{s_1}\right)\|\mathbf{\Theta}\|_1 \le \frac{m\|\widetilde{\boldsymbol{X}}\mathbf{\Theta}\|_2^2}{2} + \frac{5(K+1)q\left(\sqrt{M}\eta^*\Delta^U+\frac{1}{s_1}\right)^2}{m\|\widetilde{\boldsymbol{X}}\|_2^2\phi^2} - \left(\sqrt{M}\eta^*\Delta^U + \frac{1}{s_1}\right)\|\mathbf{\Theta}\|_1. \tag{B.3.2}$$

Next, using $\min w_i \ge m$, $\|\boldsymbol{\varepsilon}\|_\infty \lesssim \sqrt{\log ns}$ and $\|\widetilde{\boldsymbol{X}}\|_2^2 \ge ns$ and conditions (i) and (ii) in Nie and Ročková [2023], and substituting (B.3.2) in (B.3.1), we get,

$$\frac{m}{2}\|\widetilde{\boldsymbol{X}}\mathbf{\Theta}\|_2^2 + \left(\sqrt{M}\eta^*\Delta^U + \frac{1}{s_1}\right)\|\mathbf{\Theta}\|_1 < \frac{C_5^2M(\eta^*)^2}{m\phi^2}\,q(1+K)\log p.$$

Therefore, $\|\widetilde{\boldsymbol{X}}\boldsymbol{\Theta}\|_2 \leq \frac{C_5\eta^*\sqrt{M}}{\sqrt{m}\,\phi}\sqrt{q(1+K)\log p}$ and the definition of $c$ would imply

$$\lim_{n\to\infty} \mathbb{P}_{\boldsymbol{w},\boldsymbol{\mu},\boldsymbol{\beta_0}}\left(\|\boldsymbol{\Theta}\|_2 \leq \frac{C_5\eta^*\sqrt{M}}{\sqrt{m}\,\phi\, c}\sqrt{\frac{q(1+K)\log p}{ns}}\right) \geq 1. \tag{B.3.3}$$

Second, note that the difference $\widetilde{\boldsymbol{\beta}} - \hat{\boldsymbol{\beta}}$ depends on $\boldsymbol{\mu}$ and therefore, for any $z \geq 0$, applying Markov's inequality, we obtain,

$$\mathbb{P}_{\boldsymbol{\mu}}\left(\|\widetilde{\boldsymbol{\beta}} - \hat{\boldsymbol{\beta}}\|_2 > z \,\middle|\, \widetilde{\boldsymbol{y}}\right) \leq \frac{1}{z}\mathbb{E}\left[\sum_{j=1}^{d}\sum_{k=1}^{p_j}(\widetilde{\beta}_{jk} - \hat{\beta}_{jk})^2 \,\middle|\, \widetilde{\boldsymbol{y}}\right] \leq \frac{1}{z}\sum_{j=1}^{d} p_j 3 s_0^2 = \frac{3ps_0^2}{z}. \tag{B.3.4}$$

where $p = \sum_{j=1}^{d} p_j$. Now, choosing $z = \left(\frac{C_5\eta^*\sqrt{M}}{\sqrt{m}\,\phi\, c}\sqrt{\frac{q(1+K)\log p}{ns}}\right)^2$ and using (B.3.3) and (B.3.4), we obtain,

$$\lim_{n\to\infty} \mathbb{E}_{\boldsymbol{\beta}_0}\mathbb{P}_{\boldsymbol{w},\boldsymbol{\mu}}\left(\|\widetilde{\boldsymbol{\beta}} - \boldsymbol{\beta_0}\|_2 > \sqrt{z} \,\middle|\, \widetilde{\boldsymbol{y}}\right) = 0.$$

### C Additional Simulations

In this section, we present an additional simulation scenario with a smaller sample size of $n = 200$ and a fixed number of predictors $p = 658$. The goal of this setting is to evaluate the robustness of our proposed method in a more challenging, low-sample, high-dimensional regime. The data generation process follows the same setup as described in Section 3 of the main manuscript, ensuring consistency in the underlying signal structure and allowing for a meaningful comparison with previously reported results.

The simulation results reaffirm the findings reported in the main manuscript: BayesCOOP continues to demonstrate substantial improvements in both mean squared error (MSE) and mean squared prediction error (MSPE) relative to existing multiview methods (**Figure S1**-**Figure S2**). These results highlight the method's stability and effectiveness even with reduced sample sizes, and support the overall conclusion that BayesCOOP offers a powerful and scalable solution for high-dimensional multiview integration.

### D Additional Analyses

In this section, we present some additional investigations for the real datasets considered in the main manuscript. Specifically, we demonstrate the feature selection performance and the predictive uncertainty quantification of BayesCOOP, respectively for the StelzerEGA and the StelzerDOS datasets (**Figure S3**-

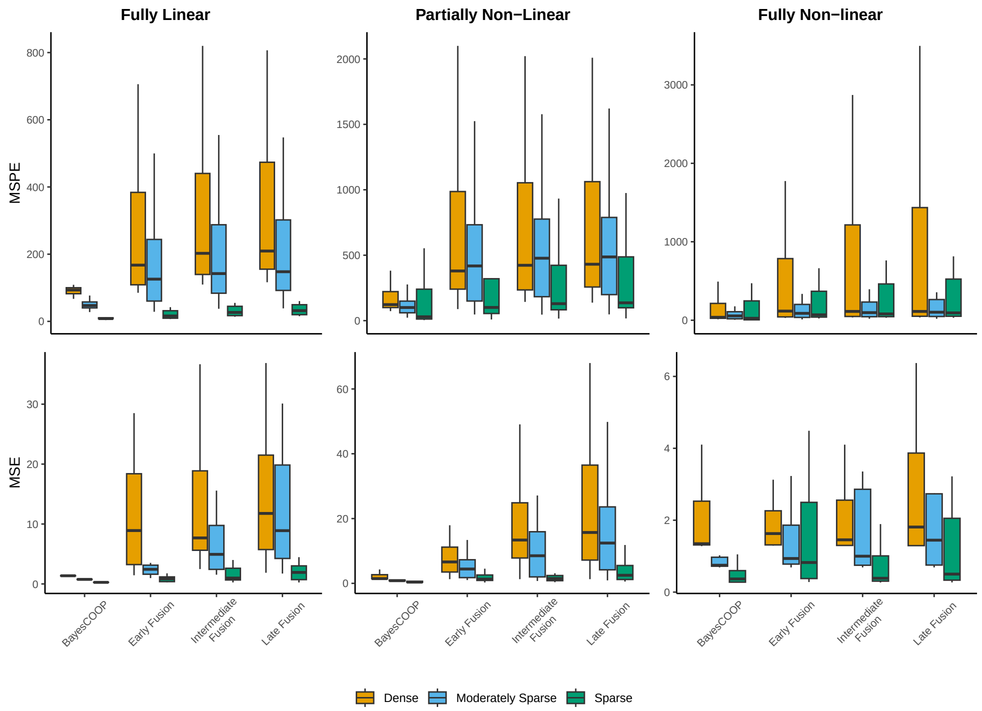


Figure S1: **BayesCOOP significantly outperformed published methods in estimation (MSE) and prediction (MSPE) in low SNR (SNR = 5) settings with sample size 200**. The values are based on completely held-out test samples, summarized over 25 iterations.

**Figure S4**).

### E Sensitivity Analysis of BayesCOOP on Real Datasets

In this section, we investigate the sensitivity of our model's performance to different choices of hyperparameters across the real datasets analyzed in the main manuscript. Specifically, we focus on the two scale parameters of the double exponential prior, denoted by $s_0$ and $s_1$.

To assess robustness, we systematically vary these hyperparameters following Yi et al. [2018] and evaluate the resulting predictive performance. As shown in **Table S1–Table S2**, the results indicate that our method exhibits strong stability across a wide range of hyperparameter configurations. This suggests that BayesCOOP is not overly sensitive to the precise specification of $s_0$ and $s_1$, making it a reliable choice in practice even when prior knowledge about appropriate scale values is limited.

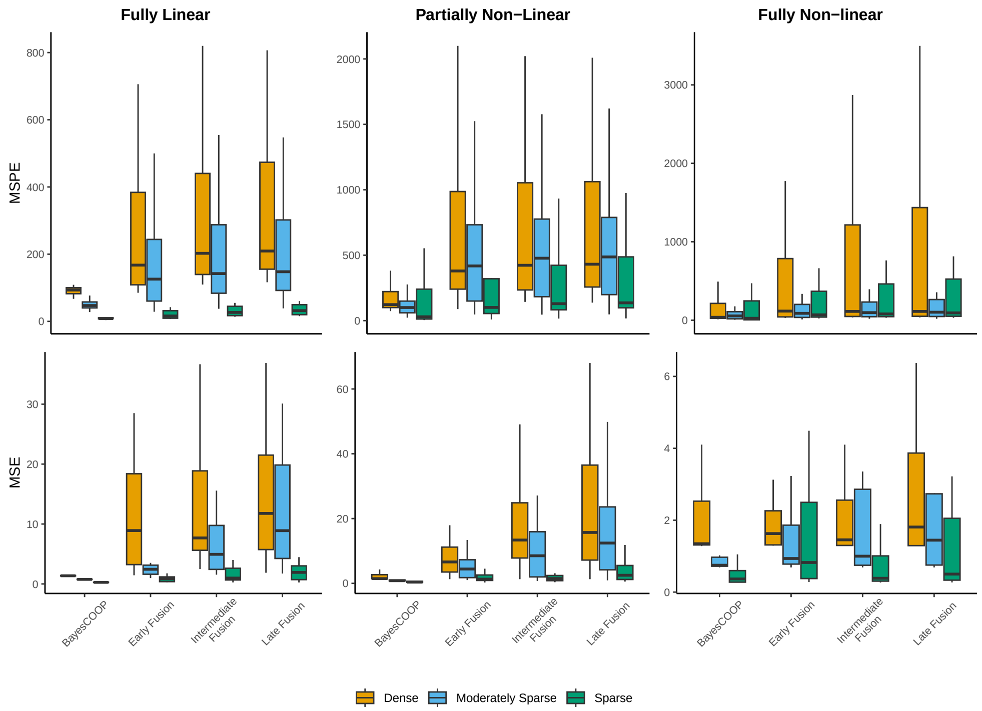


Figure S2: **BayesCOOP significantly outperformed published methods in estimation (MSE) and prediction (MSPE) in high SNR (SNR = 10) settings with sample size 200**. The values are based on completely held-out test samples, summarized over 25 iterations.

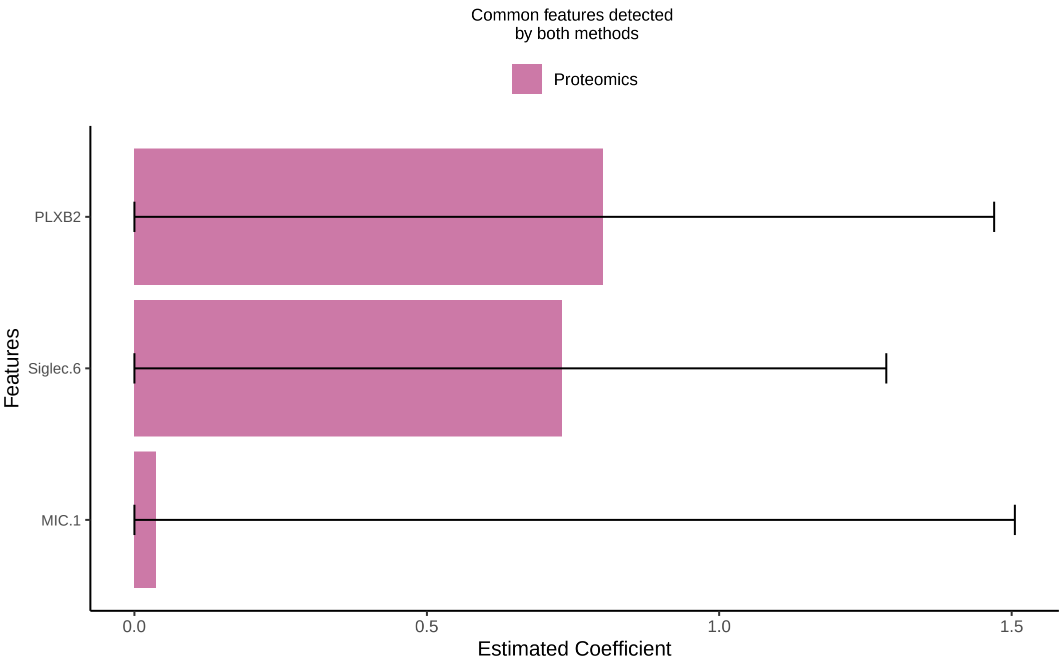


Figure S3: **BayesCOOP finds several biologically relevant candidate features in the StelzerEGA dataset [Stelzer et al., 2021]**. Effect sizes (posterior medians and credible intervals) for selected multiview features detected by both Cooperative Learning [Ding et al., 2022] and BayesCOOP are shown. No unique feature was detected exclusively by BayesCOOP in this dataset.

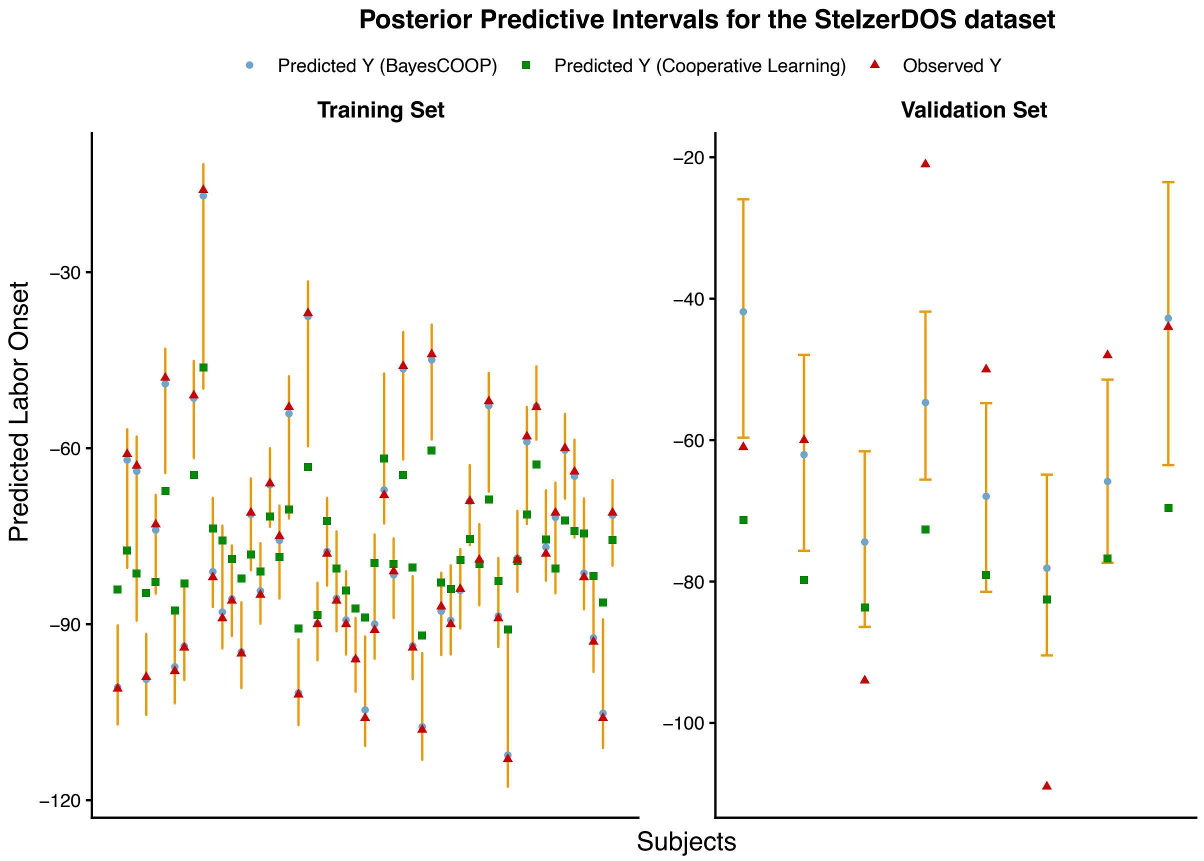


Figure S4: **BayesCOOP delivers well-calibrated and more accurate predictions of labor onset** in both the training and validation sets of the StelzerDOS dataset [Stelzer et al., 2021], outperforming Cooperative Learning [Ding et al., 2022], which provides no quantification of uncertainty in its estimates.

Table S1: Hyperparameter sensitivity analysis for the StelzerDOS dataset [Stelzer et al., 2021].

| $s_1$ | $s_0$ | MSPE | Number of features selected |
|---|---|---|---|
| 0.5 | 0.03 | 490.20 | 17 |
| | 0.05 | 494.10 | 18 |
| | 0.08 | 560.20 | 16 |
| | 0.1 | 563.93 | 15 |
| 1 | 0.03 | 489.28 | 17 |
| | 0.05 | 493.06 | 18 |
| | 0.08 | 553.09 | 14 |
| | 0.1 | 583.62 | 14 |

### F Mathematical Derivations for Algorithm 1

In this section, we present the multiview group spike-and-slab objective function and outline the mathematical derivations underlying the EM algorithm with the Bayesian bootstrap, as described in **Algorithm 1** of the main manuscript.

Table S2: Hyperparameter sensitivity analysis for the StelzerEGA dataset [Stelzer et al., 2021].

| $s_1$ | $s_0$ | MSPE | Number of features selected |
|---|---|---|---|
| 0.5 | 0.03 | 5.25 | 3 |
| | 0.05 | 7.08 | 2 |
| | 0.08 | 7.37 | 2 |
| | 0.1 | 8.52 | 7 |
| 1 | 0.03 | 5.14 | 3 |
| | 0.05 | 6.98 | 3 |
| | 0.08 | 7.15 | 2 |
| | 0.1 | 8.53 | 3 |

#### F.1 The multiview group spike-and-slab objective function

Define $\boldsymbol{\gamma} = (\gamma_{11}, \ldots, \gamma_{d,p_d})^\top$ and $\boldsymbol{\delta} = (\delta_1, \ldots, \delta_d)^\top$ and $\widetilde{\boldsymbol{X}} = (\widetilde{\boldsymbol{X}}_1, \ldots, \widetilde{\boldsymbol{X}}_{ns})^\top$. Our implementation of the EM algorithm is based on the log joint posterior of the parameters $\boldsymbol{\Lambda} = (\boldsymbol{\beta}, \sigma^2, \boldsymbol{\gamma}, \boldsymbol{\delta})$ given by,

$$
\begin{aligned}
\log\ \pi(\boldsymbol{\Lambda}|\widetilde{\boldsymbol{y}}, \widetilde{\boldsymbol{X}}) &= \sum_{i=1}^{ns} w_i \log\ \pi(\widetilde{y}_i|\widetilde{\boldsymbol{X}}_i\boldsymbol{\beta}, \sigma^2) + \sum_{j=1}^{d}\sum_{k=1}^{p_j} \log\ \pi(\beta_{jk}|\gamma_{jk}) \\
&+ \sum_{u=1}^{d}\sum_{j=1}^{d}\sum_{k=1}^{p_j} \log\ \pi(\gamma_{jk}|\delta_u) + \sum_{u=1}^{d} \log\ \pi(\delta_u) + \log\ \pi(\sigma^2) \\
&\propto -\sum_{i=1}^{ns} \frac{w_i(\widetilde{y}_i - \widetilde{\boldsymbol{X}}_i\boldsymbol{\beta})^\top(\widetilde{y}_i - \widetilde{\boldsymbol{X}}_i\boldsymbol{\beta})}{2\sigma^2} - \frac{(ns+\nu+2)\log\ \sigma^2}{2} \\
&- \sum_{j=1}^{d}\sum_{k=1}^{p_j} \frac{|\beta_{jk} - \mu_{jk}|}{(1-\gamma_{jk})s_0 + \gamma_{jk}s_1} + \sum_{j=1}^{d}\sum_{k=1}^{p_j} \log\ \{\delta_j^{\gamma_{jk}}(1-\delta_j)^{1-\gamma_{jk}}\} - \frac{\nu}{2\sigma^2}.
\end{aligned}
\tag{F.1.1}
$$

Treating the indicator variable $\boldsymbol{\gamma}$ as "missing data," the EM algorithm indirectly maximizes $p(\boldsymbol{\beta}, \boldsymbol{\delta}, \sigma^2|\widetilde{\boldsymbol{y}}, \widetilde{\boldsymbol{X}})$ by iteratively optimizing an objective function defined as

$$
\begin{aligned}
H\left(\boldsymbol{\beta}, \boldsymbol{\delta}, \sigma^2 \mid \boldsymbol{\beta}^{(t,k-1)}, \boldsymbol{\delta}^{(t,k-1)}, \sigma^{2(t,k-1)}\right) &= \mathbb{E}_{\boldsymbol{\gamma}|-}\left[\log\ \pi(\boldsymbol{\Lambda}|\widetilde{\boldsymbol{y}}, \widetilde{\boldsymbol{X}}, \boldsymbol{\beta}^{(t,k-1)}, \boldsymbol{\delta}^{(t,k-1)}, \sigma^{2(t,k-1)})\right] \\
&= C + H_1\left(\boldsymbol{\beta}, \sigma^2 \mid \boldsymbol{\beta}^{(t,k-1)}, \boldsymbol{\delta}^{(t,k-1)}, \sigma^{2(t,k-1)}\right) \\
&\quad + H_2\left(\boldsymbol{\delta} \mid \boldsymbol{\beta}^{(t,k-1)}, \boldsymbol{\delta}^{(t,k-1)}, \sigma^{2(t,k-1)}\right),
\end{aligned}
\tag{F.1.2}
$$

where $\mathbb{E}_{\gamma|-}(\cdot)$ is the conditional expectation $\mathbb{E}_{\gamma|\widetilde{\boldsymbol{y}},\widetilde{\boldsymbol{X}},\boldsymbol{\beta}^{(t,k)},\boldsymbol{\delta}^{(t,k)},\sigma^{2(t,k)}}(\cdot)$ and

$$\begin{aligned}
H_1\left(\boldsymbol{\beta},\sigma^2 \mid \boldsymbol{\beta}^{(t,k-1)},\boldsymbol{\delta}^{(t,k-1)},\sigma^{2(t,k-1)}\right) &= -\sum_{i=1}^{ns}\frac{w_i(\widetilde{y}_i-\widetilde{\boldsymbol{X}}_i\boldsymbol{\beta})^\top(\widetilde{y}_i-\widetilde{\boldsymbol{X}}_i\boldsymbol{\beta})}{2\sigma^2}-\frac{(ns+\nu+2)\log\ \sigma^2}{2}\\
&\quad -\sum_{j=1}^{d}\sum_{k=1}^{p_j}|\beta_{jk}-\mu_{jk}|\ \mathbb{E}_{\gamma|-}\left[\frac{1}{s_0(1-\gamma_{jk})+s_1\gamma_{jk}}\right]-\frac{\nu}{2\sigma^2},\\
H_2\left(\boldsymbol{\delta}\mid \boldsymbol{\beta}^{(t,k-1)},\boldsymbol{\delta}^{(t,k-1)},\sigma^{2(t,k-1)}\right) &= \sum_{j=1}^{d}\sum_{k=1}^{p_j}\mathbb{E}_{\gamma|-}[\gamma_{jk}]\log\ \left(\frac{\delta_j}{1-\delta_j}\right)+\sum_{j=1}^{d}p_j\log\ (1-\delta_j).
\end{aligned}\tag{F.1.3}$$

At $k^{\text{th}}$ iteration, starting from the current estimates $\boldsymbol{\beta}^{(t,k-1)},\boldsymbol{\delta}^{(t,k-1)},\sigma^{2(t,k-1)}$, the algorithm calculates the expectation of the expression on the right side of (F.1.2), resulting in the function $H$. Maximizing $H$ with respect to $\boldsymbol{\beta},\boldsymbol{\delta},\sigma^2$ yields the updated values $(\boldsymbol{\beta}^{(t,k)},\boldsymbol{\delta}^{(t,k)},\sigma^{2(t,k)})$.

#### F.2 The EM Coordinate Descent Algorithm

Let us now describe the EM algorithm for the pseudo-MAP estimation of the high-dimensional regression coefficient $\boldsymbol{\beta}$ under each Bayesian bootstrap draw. The following sections describe these two steps in detail.

**E-step** The E-step proceeds by computing the terms $\mathbb{E}_{\gamma|-}[\gamma_{jk}]$ and $\mathbb{E}_{\gamma|-}\left[\frac{1}{s_0(1-\gamma_{jk})+s_1\gamma_{jk}}\right]$ in (F.1.3). Clearly,

$$\mathbb{E}_{\gamma|-}[\gamma_{jk}]=\pi(\gamma_{jk}=1\mid \boldsymbol{\beta}^{(t,k-1)},\boldsymbol{\delta}^{(t,k-1)},\sigma^{2(t,k-1)})=\frac{u_{jk}}{u_{jk}+v_{jk}},$$

where $u_{jk}=\pi(\beta_{jk}\mid\gamma_{jk}=1,s_1)\pi(\gamma_{jk}=1\mid\delta_j)$, $v_{jk}=\pi(\beta_{jk}\mid\gamma_{jk}=0,s_0)\pi(\gamma_{jk}=0\mid\delta_j)$ and

$$\mathbb{E}_{\gamma|-}\left[\frac{1}{s_0(1-\gamma_{jk})+s_1\gamma_{jk}}\right]=\frac{1}{u_{jk}+v_{jk}}\left(\frac{v_{jk}}{s_0}+\frac{u_{jk}}{s_1}\right).$$

Substituting these values into (F.1.2) provides the objective function $H(\cdot)$ in analytical form, which we maximize in the next step.

**M-step** The decomposition of the objective function $H\left(\boldsymbol{\beta},\boldsymbol{\delta},\sigma^2\mid\boldsymbol{\beta}^{(t,k-1)},\boldsymbol{\delta}^{(t,k-1)},\sigma^{2(t,k-1)}\right)$ into two independent components simplifies the maximization process by allowing each component to be optimized separately, significantly enhancing the efficiency of the M-step in the algorithm: $H_1\left(\boldsymbol{\beta},\sigma^2\mid\boldsymbol{\beta}^{(t,k-1)},\boldsymbol{\delta}^{(t,k-1)},\sigma^{2(t,k-1)}\right)$ and $H_2\left(\boldsymbol{\delta}\mid\boldsymbol{\beta}^{(t,k-1)},\boldsymbol{\delta}^{(t,k-1)},\sigma^{2(t,k-1)}\right)$.

In the first part of the M-step, $H_1(\boldsymbol{\beta}, \sigma^2)$ is maximized. This process is carried out in two stages. First, regardless of the value of $\sigma^{2(t,k)}$, the parameter $\boldsymbol{\beta}^{(t,k)}$ is obtained by solving a weighted LASSO regression problem using a fast cyclic coordinate descent algorithm [Friedman et al., 2010; Tang et al., 2018]. Specifically, $\boldsymbol{\beta}^{(t,k)}$ is updated as

$$\boldsymbol{\beta}^{(t,k)} = \arg\min_{\boldsymbol{\beta}\in\mathbb{R}^p} \sum_{i=1}^{ns} w_i(\widetilde{y}_i - \widetilde{\boldsymbol{X}}_i\boldsymbol{\beta})^\top(\widetilde{y}_i - \widetilde{\boldsymbol{X}}_i\boldsymbol{\beta}) + \sum_{j=1}^{d}\sum_{k=1}^{p_j} \lambda_{jk} \left|\beta_{jk} - \mu_{jk}\right|,$$

where the penalty parameter $\lambda_{jk} = \frac{1}{u_{jk}+v_{jk}}\left(\frac{v_{jk}}{s_0} + \frac{u_{jk}}{s_1}\right)$. This step enables automatic variable selection, as some coefficients of $\boldsymbol{\beta}$ naturally reduce to exactly zero. Once $\boldsymbol{\beta}^{(t,k)}$ is determined, the value of $\sigma^{2(t,k)}$ is updated using the following formula:

$$\sigma^{2(t,k)} = \frac{\sum_{i=1}^{ns} w_i(\widetilde{y}_i - \widetilde{\boldsymbol{X}}_i\boldsymbol{\beta}^{(t,k)})^\top(\widetilde{y}_i - \widetilde{\boldsymbol{X}}_i\boldsymbol{\beta}^{(t,k)}) + \nu}{ns + \nu + 2}.$$

In the second part of the M-step, $H_2(\boldsymbol{\delta})$ is maximized with respect to $\boldsymbol{\delta}$. The parameter $\boldsymbol{\delta}^{(t,k)}$ is obtained as a closed-form solution to the following expression:

$$\delta_j^{(t,k)} = \arg\max_{\delta\in\mathbb{R}} \sum_{k=1}^{p_j} \frac{u_{jk}}{u_{jk}+v_{jk}} \log\left(\frac{\delta_j}{1-\delta_j}\right) + p_j \log\,(1-\delta_j),$$

which is given by $\delta_j^{(t,k)} = \frac{1}{p_j}\sum_{k=1}^{p_j}\frac{u_{jk}}{u_{jk}+v_{jk}}$. The convergence of the EM algorithm is assessed using the criterion $\frac{|d^{(t,k)}-d^{(t,k-1)}|}{0.1+|d^{(t,k)}|} < \epsilon$, where $d^{(t,k)} = -2\sum_{i=1}^{ns}\log\,\pi(\widetilde{y}_i \mid \widetilde{\boldsymbol{X}}_i\boldsymbol{\beta}, \sigma^2)$ represents the deviance estimate at the $k^{\text{th}}$ iteration, and $\epsilon$ is a small threshold value (e.g., $10^{-5}$).

#### F.3 Posterior Distributions

In this section, we derive the full conditional posterior distributions for the parameters $\sqrt{\rho}$ and $\sigma^2$ under the proposed Bayesian hierarchical model.

**Conditional posterior distribution of $\sqrt{\rho}$.** The conditional posterior of $\sqrt{\rho}$ is given by

$$\begin{aligned} f(\sqrt{\rho} \mid \widetilde{\boldsymbol{y}}, \boldsymbol{\beta}, \sigma^2) &\propto f(\widetilde{\boldsymbol{y}} \mid \boldsymbol{\beta}, \rho, \sigma^2) \cdot f(\sqrt{\rho}) \\ &\propto \exp\left(-\frac{\|\widetilde{\boldsymbol{y}} - \widetilde{\boldsymbol{X}}\boldsymbol{\beta}\|_2^2}{2\sigma^2}\right) \cdot \mathbb{I}_{(0,1)}(\sqrt{\rho}) \\ &\propto \exp\left(-\frac{(\sqrt{\rho})^2 \sum_{j<k} \left\|\boldsymbol{X}^{(j)}\boldsymbol{\beta}_j - \boldsymbol{X}^{(k)}\boldsymbol{\beta}_k\right\|_2^2}{2\sigma^2}\right) \cdot \mathbb{I}_{(0,1)}(\sqrt{\rho}). \end{aligned}$$

This expression is proportional to the kernel of a truncated normal distribution, leading to

$$\sqrt{\rho} \mid \widetilde{\boldsymbol{y}}, \boldsymbol{\beta}, \sigma^2 \sim \mathcal{N}\left(0, \frac{\sigma^2}{\sum_{j<k} \|\boldsymbol{X}^{(j)}\boldsymbol{\beta}_j - \boldsymbol{X}^{(k)}\boldsymbol{\beta}_k\|_2^2}\right) \cdot \mathbb{I}_{(0,1)}(\sqrt{\rho}).$$

**Conditional posterior distribution of $\sigma^2$.** Similarly, the conditional posterior of $\sigma^2$ can be derived as

$$\begin{aligned} f(\sigma^2 \mid \widetilde{\boldsymbol{y}}, \boldsymbol{\beta}, \rho) &\propto f(\widetilde{\boldsymbol{y}} \mid \boldsymbol{\beta}, \rho, \sigma^2) \cdot f(\sigma^2) \\ &\propto (\sigma^2)^{-(ns+\nu)/2} \exp\left(-\frac{\nu + \sum_{i=1}^{ns} \|\widetilde{y}_i - \widetilde{\boldsymbol{X}}_i\boldsymbol{\beta}\|_2^2}{2\sigma^2}\right), \end{aligned}$$

which corresponds to an inverse-gamma distribution:

$$\sigma^2 \mid \widetilde{\boldsymbol{y}}, \boldsymbol{\beta}, \rho \sim \text{Inverse-Gamma}\left(\frac{ns+\nu}{2}, \frac{\nu + \sum_{i=1}^{ns} \|\widetilde{y}_i - \widetilde{\boldsymbol{X}}_i\boldsymbol{\beta}\|_2^2}{2}\right).$$

### References


Albert, J. H. and Chib, S. (1993). Bayesian analysis of binary and polychotomous response data. *Journal of the American Statistical Association*, 88(422):669–679.

Ding, D. Y., Li, S., Narasimhan, B., and Tibshirani, R. (2022). Cooperative learning for multiview analysis. *Proceedings of the National Academy of Sciences*, 119(38):e2202113119.

Friedman, J. H., Hastie, T. J., and Tibshirani, R. (2010). A note on the group lasso and a sparse group lasso. *arXiv: Statistics Theory*.

Kleinbaum, D. G. and Klein, M. (2006). *Survival Analysis: A Self-Learning Text.* Springer Science and Business Media, LLC, New, 2nd edition.

Leng, C., Tran, M.-N., and Nott, D. (2014). Bayesian adaptive lasso. *Annals of the Institute of Statistical Mathematics*, 66(2):221–244. Received July 4, 2012; Revised May 6, 2013; Published September 3, 2013.

Maity, A. K., Bhattacharya, A., Mallick, B. K., and Baladandayuthapani, V. (2019). Bayesian data integration and variable selection for pan-cancer survival prediction using protein expression data. *Biometrics*, 76(1):316–325.

Mallick, H., Porwal, A., Saha, S., Basak, P., Svetnik, V., and Paul, E. (2024). An integrated bayesian framework for multi-omics prediction and classification. *Statistics in Medicine*, 43(5):983–1002.

Nie, L. and Ročková, V. (2023). Bayesian bootstrap spike-and-slab lasso. *Journal of the American Statistical Association*, 118(543):2013–2028.

Ročková, V. and George, E. I. (2018). The spike-and-slab lasso. *Journal of the American Statistical Association*, 113(521):431–444.

Sha, N., Tadesse, M. G., and Vannucci, M. (2006). Bayesian variable selection for the analysis of microarray data with censored outcomes. *Bioinformatics*, 22(18):2262–2268.

Stelzer, I. A. et al. (2021). Integrated trajectories of the maternal metabolome, proteome, and immunome predict labor onset. *Science Translational Medicine*, 13(592).

Tang, Z., Shen, Y., Li, Y., Zhang, X., Wen, J., Qian, C., Zhuang, W., Shi, X., and Yi, N. (2018). Group spike-and-slab lasso generalized linear models for disease prediction and associated genes detection by incorporating pathway information. *Bioinformatics*, 34(6):901–910.

Wang, H. and Leng, C. (2007). Unified lasso estimation by least squares approximation. *Journal of the American Statistical Association*, 102(479):1039–1048.

Yi, N., Tang, Z., Zhang, X., and Guo, B. (2018). Bhglm: Bayesian hierarchical glms and survival models, with applications to genomics and epidemiology. *Bioinformatics*, 35(8):1419–1421.
